# A dinoflagellate-infecting giant virus with a micron-length tail

**DOI:** 10.1101/2025.07.19.665647

**Authors:** Andrian P. Gajigan, Christopher R. Schvarcz, Amanda B. Laughlin, Tina M. Weatherby, Alexander I. Culley, Kyle F. Edwards, Grieg F. Steward

## Abstract

Viral infection is a ubiquitous source of marine plankton mortality, but relatively few viruses that infect phytoplankton have been characterized. Here we describe a virus, PelV-1, with unusual morphological and genomic features that infects a dinoflagellate, *Pelagodinium* sp. Both host and virus were isolated from the epipelagic zone in the North Pacific Subtropical Gyre. PelV-1 has a ∼200 nm capsid size, and the virion variably exhibits two appendages, the presence and length of which may reflect different stages of virion maturity or artifacts of sample preparation. The appendages are a thinner 30 nm-wide tail-like structure that can extend to 2.3 µm — the longest virus appendage described to date— and a shorter, thicker (>40–70 nm) protrusion, which appears to emerge from a star-shaped capsid opening directly opposite the attachment point of the long, thin tail. Sequencing and assembly of material in a purified lysate generated a high-coverage (> 4,000×) genome of 459 kb (33.8% GC). A second, distinct genome of 504 kb (25.8% GC) was also assembled, but had low read coverage (< 24×), suggesting the presence of a low-abundance, co-cultured virus (co-PelV). Phylogenetic analysis indicates that both PelV-1 and co-PelV are members of *Mesomimiviridae*. They contain various genes for the metabolism of amino acids (e.g., asparagine synthase), carbohydrates (e.g., epimerase, glycosyl hydrolase, aconitate hydratase, succinate dehydrogenase of the TCA cycle), and lipids (e.g., phospholipases), as well as other noteworthy genes (e.g., light-harvesting complex, rhodopsin, ion channel, sugar transporters, aquaporin). PelV-1 also has ORFs most similar to tail fiber genes of *Synechococcus* phage and other tail domain-containing protein homologs. The ecological advantages that might be conferred by the extraordinarily long tail and metabolic genes of PelV-1 is unknown, but this isolate expands the scope of morphological and metabolic diversity of viruses and suggests many more unusual marine viruses await discovery.

**Author summary:** Giant viruses challenged our traditional views of virology due to their large size and the presence of hundreds of auxiliary metabolic genes. But despite the immense giant virus diversity discovered through sequencing, few isolates were described, and those were primarily viruses that infect amoeba host and rarely from phytoplankton. This hampers our understanding of marine host-virus interaction and thus the impact of viruses on the ocean ecosystem. Here we provide genomic and morphological characterization of a novel dinoflagellate giant virus (PelV-1) and a second co-occurring, albeit low abundance, virus (co-PelV). Dinoflagellates are vital in marine symbiosis and algal blooms but only two giant virus isolates have been described with no available genomic resources to date. Thus, this is a significant contribution to the literature on dinoflagellate viruses. Among the notable features of PelV-1 are its unique micron-length tail appendage, phagocytosis-like entry mechanism and its varied auxiliary metabolic genes including photosynthesis and energy generating genes.

## Introduction

The characterization of the largest, or “giant”, virus isolates in the phylum *Nucleocytoviricota* has led to a rapid expansion of the known universe of genes encoded within the virosphere, including diverse collections of functional genes involved in protein translation (Abrahão et al., 2018), lipid metabolism (Blanc-Mathieu et al., 2021), tricarboxylic acid cycle, glycolysis, and gluconeogenesis (Aherfi et al., 2022; Ha et al., 2021; Moniruzzaman et al., 2020), fermentation (Schvarcz & Steward, 2018), photosynthesis (Needham et al., 2019; Schulz et al., 2020), and ion transport and assimilation (Monier et al., 2017; Moniruzzaman et al., 2020), among others. Despite the notable phylogenetic diversity among giant virus isolates described to date, it is clear we have still just scratched the surface. An extensive multi-environment and global sequencing survey, for example, showed that we lack cultured isolates for most groups of giant viruses (Schulz et al., 2020). Concerted efforts to isolate and characterize more viruses across a broad spectrum of eukaryotic hosts would add value to such environmental sequencing efforts, by better illuminating the relationship between virus genotype and phenotype and deepening our understanding of the virus-host interactions.

One significant group of eukaryotes for which there are still few characterized viruses is the dinoflagellates (Short et al., 2020). Dinoflagellates (class Dinophyceae) are a diverse group in terms of their modes of nutrition (autotrophic, heterotrophic, or mixotrophic), habitats (freshwater, estuarine, marine, and even in snow or ice) (Hoham & Remias, 2020), species richness, and lifestyles (Durvasula, 2020). Dinoflagellates are frequent partners in marine symbioses, major contributors to the microbial food web as both primary producers and grazers, and as causative species of harmful algal blooms, with significant ramifications for public health and economies, but how viruses influence dinoflagellate ecology remains little known. As early as 1979, large (385 ± 5 nm) icosahedral virus-like particles (VLPs) were observed by electron microscopy in dinoflagellate *Gymnodinium uberrimum* (Sicko-Goad & Walker, 1979), but the particles were never isolated or further studied. Filamentous VLPs have also been observed in the coral symbionts in the genus *Symbiodinium* (Howe-Kerr et al., 2023) and sequencing suggest dsDNA and ssRNA viruses may be present in such systems (Correa et al., 2013; Grupstra et al., 2022; S. A. Lawrence et al., 2017), but these have also not been cultivated. Among the known isolates of dinoflagellate-infecting viruses are a suite of small RNA viruses (HcRNAV) that infect strains of the bivalve-killing *Heterocapsa circularisquama* (Nagasaki et al., 2005; Tomaru et al., 2004). Only two other dinoflagellate-infecting virus isolates have been described, both large, double-stranded DNA viruses within the phylum *Nucleocytoviricota* that also infect members of the genus *Heterocapsa*. The first is HcDNAV, which infects *H. circularisquama* (Tarutani et al., 2001) and the second is HpygDNAV, which infects *H. pygmea* (Kim et al., 2012). A genome sequence has not been reported for either virus. Only the PolB and MutS sequences are available for HcDNAV (Ogata et al., 2009), which surprisingly placed HcDNAV as a sister clade to *Asfaviridae*, not under the common algal virus orders *Imitervirales* or *Algavirales*. With so few isolates described to date, it is evident that there is much still to learn about the diversity of viruses infecting this ubiquitous, diverse, and consequential group of protists.

Here we describe the unusual morphology, and distinctive genome features of a new virus isolate, PelV-1, that infects an open-ocean strain of dinoflagellate in the genus *Pelagodinium*. We also present a draft genome of a second virus, co-PelV, that was discovered as a minor co-occurring virus during the PelV-1 genome assembly process. PelV-1 provides a cultivated representative of a tailed giant virus that should be useful in experimentally testing the function and ecological advantages of such structures.

## Results

### Virion structural features

Virions with a capsid diameter of ∼180–200 nm and variable presence of appendages were observed in crude lysates and in equilibrium buoyant density gradient fractions in the range of 1.38–1.55 g mL^-1^ in CsCl (**S1 Fig**) or 1.16–1.18 g mL^-1^ in iodixanol (**S2 Fig**). Five different morphotypes were ascribed based on the presence and nature of two distinct appendages. Morphotype 1 contains a thin (∼30 nm width), long tail **(Fig 1A)**. Morphotype 2, the dominant type in the viral fractions **(Fig 1B, S1 Fig, S2 Fig)** has a shorter, but variable length tail of similar width (∼30 nm, though it can reach thickness up to ∼35-40 nm). Morphotype 3 has a tapered, stubby protrusion that is somewhat thicker at the base (40 to ∼70 nm width). This protrusion appears to emanate from the capsid apex, opposite the vertex where the thin tail is attached as shown in some of the negative staining **(Figs. 1C& 1F)**. We also observed a non-tailed variant, which we named morphotype 4 **(Fig. 1D)**. Lastly, we observed a virion containing both a long, thin tail (type 1 or 2) and stubby protrusion (type 3), which we labeled as morphotype 5 (**Figs. 1E& 1F)**.

**Figure 1.**
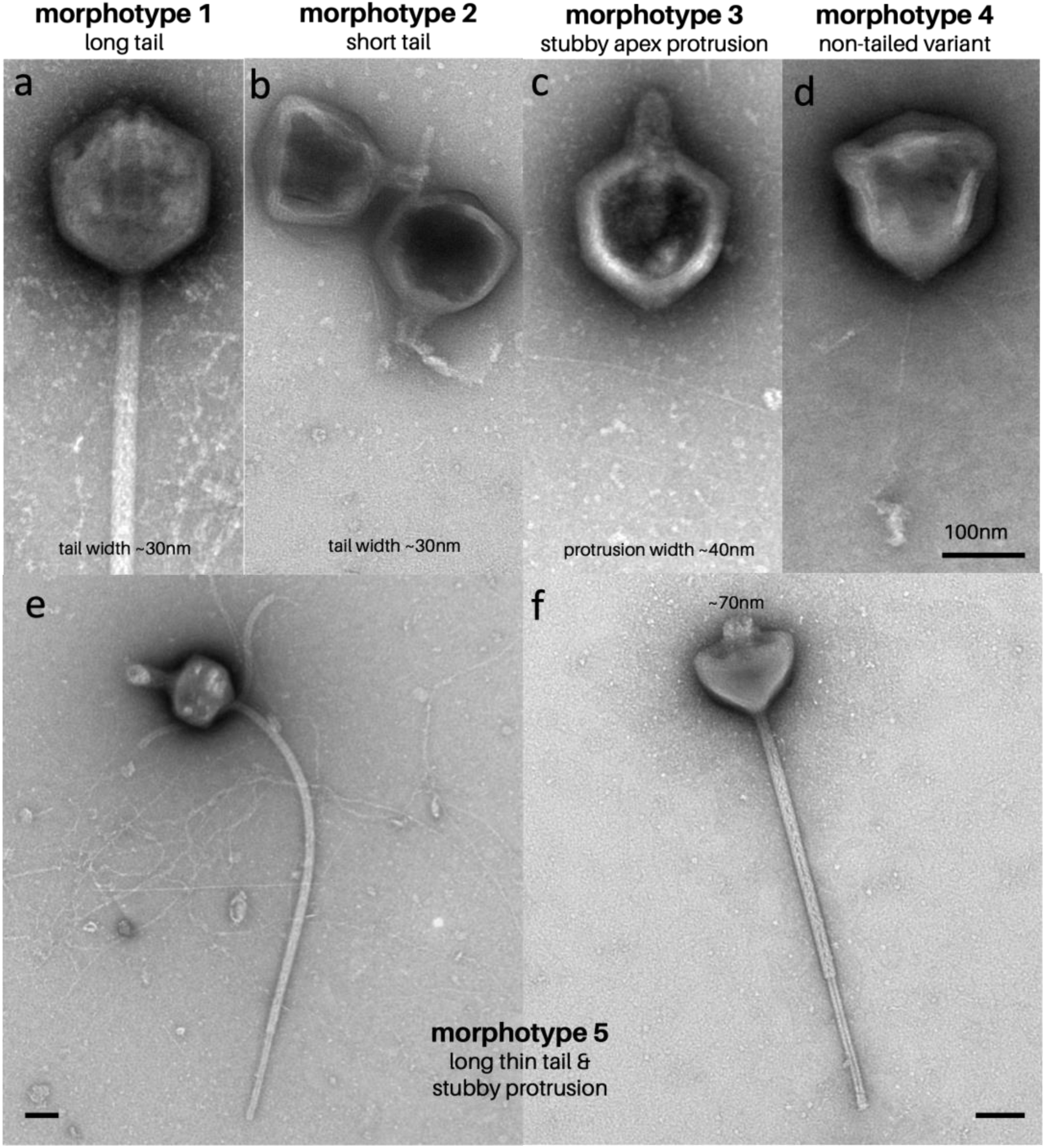
Varied morphotypes in lysate and purified fractions. (a-f) Five distinct morphotypes are observable in the lysate and fractions. **(a)** First, the long-tailed morphotype 1 is present in lysate and fractions, while the **(b)** short-tailed morphotype 2 is present in lysate and dominant in purified fractions, likely representing a broken tail or still developing tail. **(c)** We differentiate morphotype 3 from the other since it is likely related to the protrusion coming out of the apex **(e, f**). **(d)** Non-tailed variant, morphotype 4, is also present in lysate and during egress. Capsid size is between 180-200 nm **(e and f)** Morphotype 5 contains both long tail and (stubby apex protrusion). **(f)** Virion with collapsed apex, showing the stubby protrusion. Scale at 100 nm.

In undisturbed fresh lysate the long, thin tail-like structure (morphotype 1) can be observed comprising of an outer (30 nm) and inner (25 nm) tube **(Fig. 2, S3 Fig)**. In some instances, the inner tube is detached, and three putative tail fibers come out from one side **(S3 Fig)**. Similar to other giant viruses, those in our isolates contain a five-fold symmetrical “stargate” opening at the capsid apex **(Fig. 2)**. In both lysates and purified fractions, we occasionally observed filamentous structures near the capsid opening, which we call a nucleoprotein-like assembly (NLA) **(S4 Fig)**. The width and surface protein projections of the nucleoprotein-like assembly (∼16nm) are strikingly different from the virus tails (30 nm) **(Fig. S5)**.

**Figure 2.**
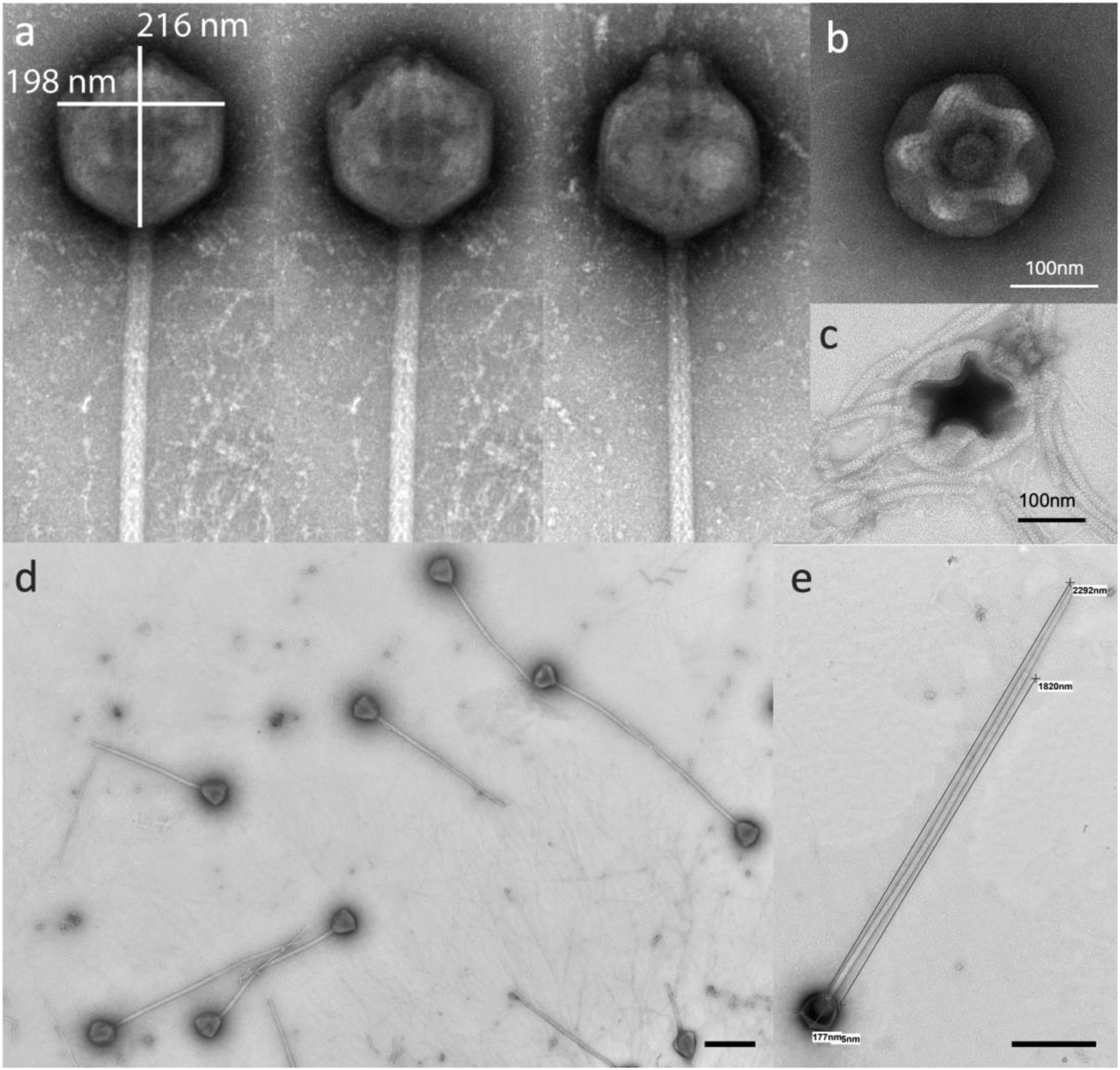
PelV-1 stargate and tail features. (a-c) Five-fold symmetry opening in the apex similar to other Mimiviruses. (d-e) Long tail PelV-1 morphotype can extend up to 2.3 microns. Scale bar is 400 nm.

### Virus attachment, entry, and egress observations from time series infection

We performed a time series infection twice to observe the virus attachment, entry, and egress **(S6 Fig & S7 Fig)**. Typical organelles are present for healthy, uninfected host cells, including the unusual dinoflagellate nucleus containing a condensed form of chromosome **(S8 Fig**). In the early stages of the infection (1, 15 mpi, and 6 hpi), virions were observed attached to the cell surface via a short thin tail **(Fig. 3)**. There was no evidence from TEM sections that the tail penetrates the cell wall **(Fig. 3)**. Entry appeared to be by endocytosis as observed from 15 mpi to 6 hpi **(Fig. 4)**. At the start of lysis (∼12 hpi), we saw no evidence of tails on the cell-associated virions observed by either SEM **(Figure 5)** or TEM **(panel C S6 Fig)**.

**Figure 3.**
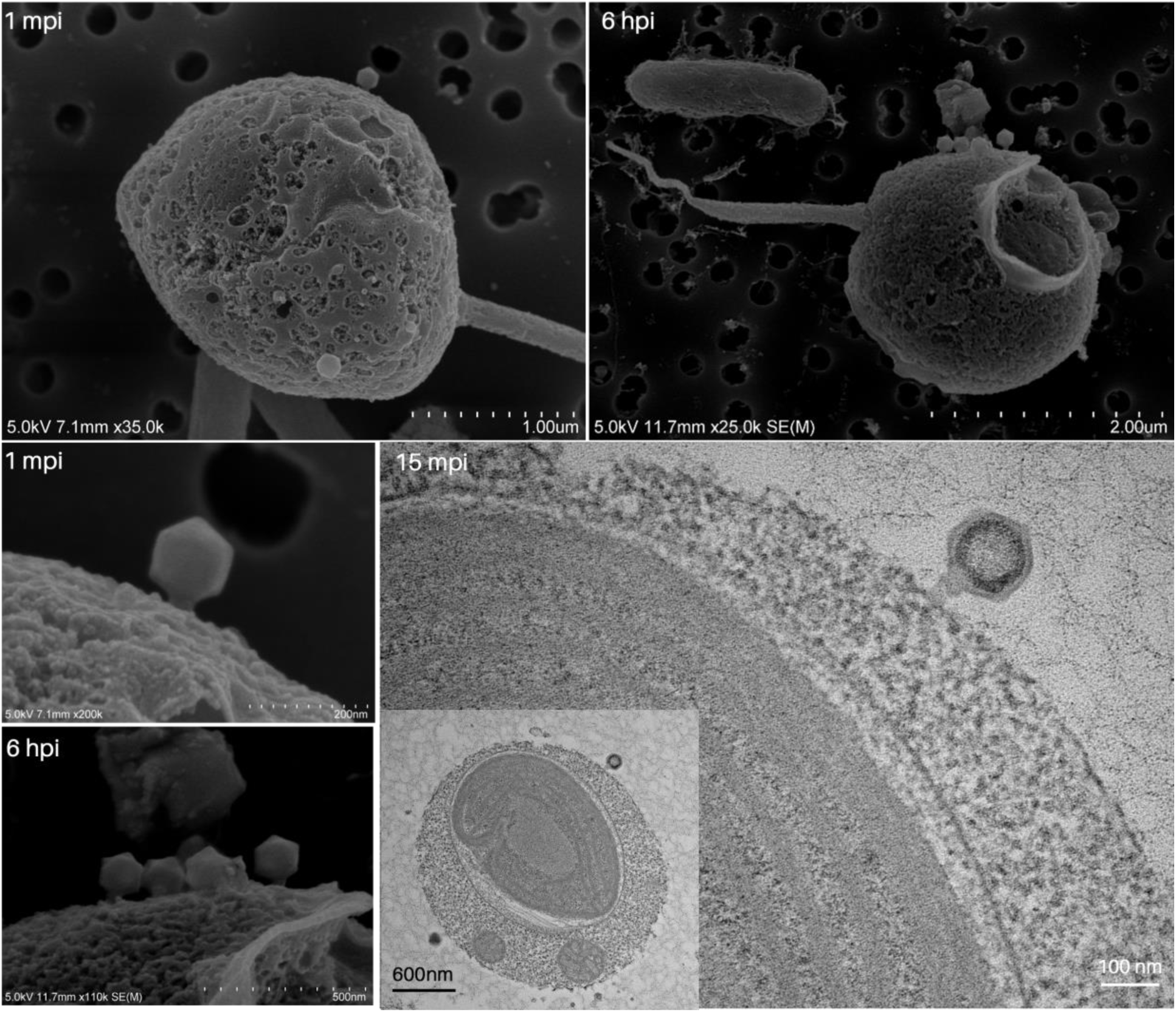
Tail attachment. Scanning (SEM) and transmission (TEM) electron micrographs of the *Pelagodinium*-PelV1 system at the early stages of infection.

**Figure 4.**
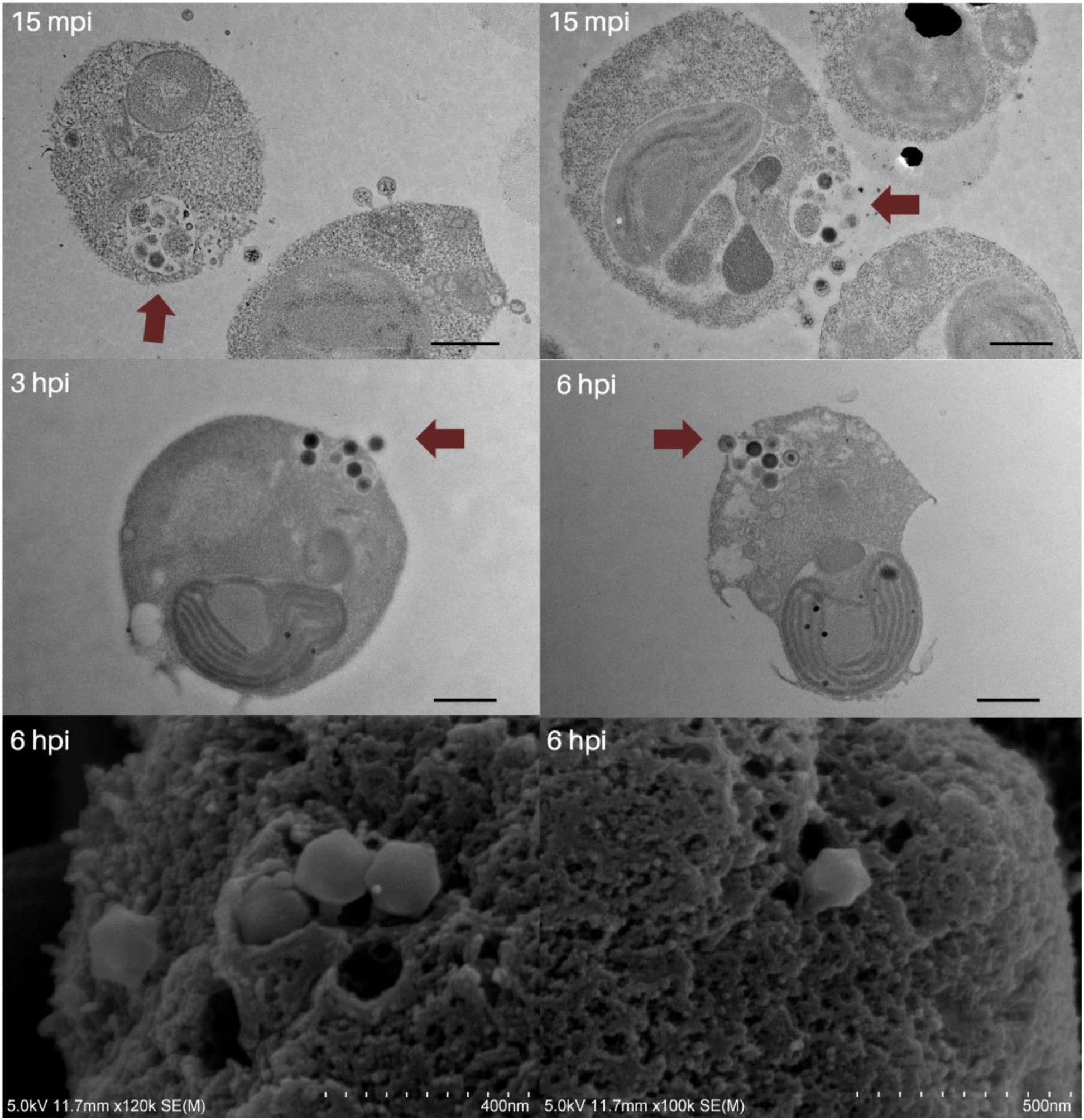
Endocytosis-like entry mechanism. Transmission and scanning electron micrographs of the *Pelagodinium*-PelV1 system at the early stages of infection show an endocytosis-like entry mechanism—scale bar at 600 nm.

**Figure 5.**
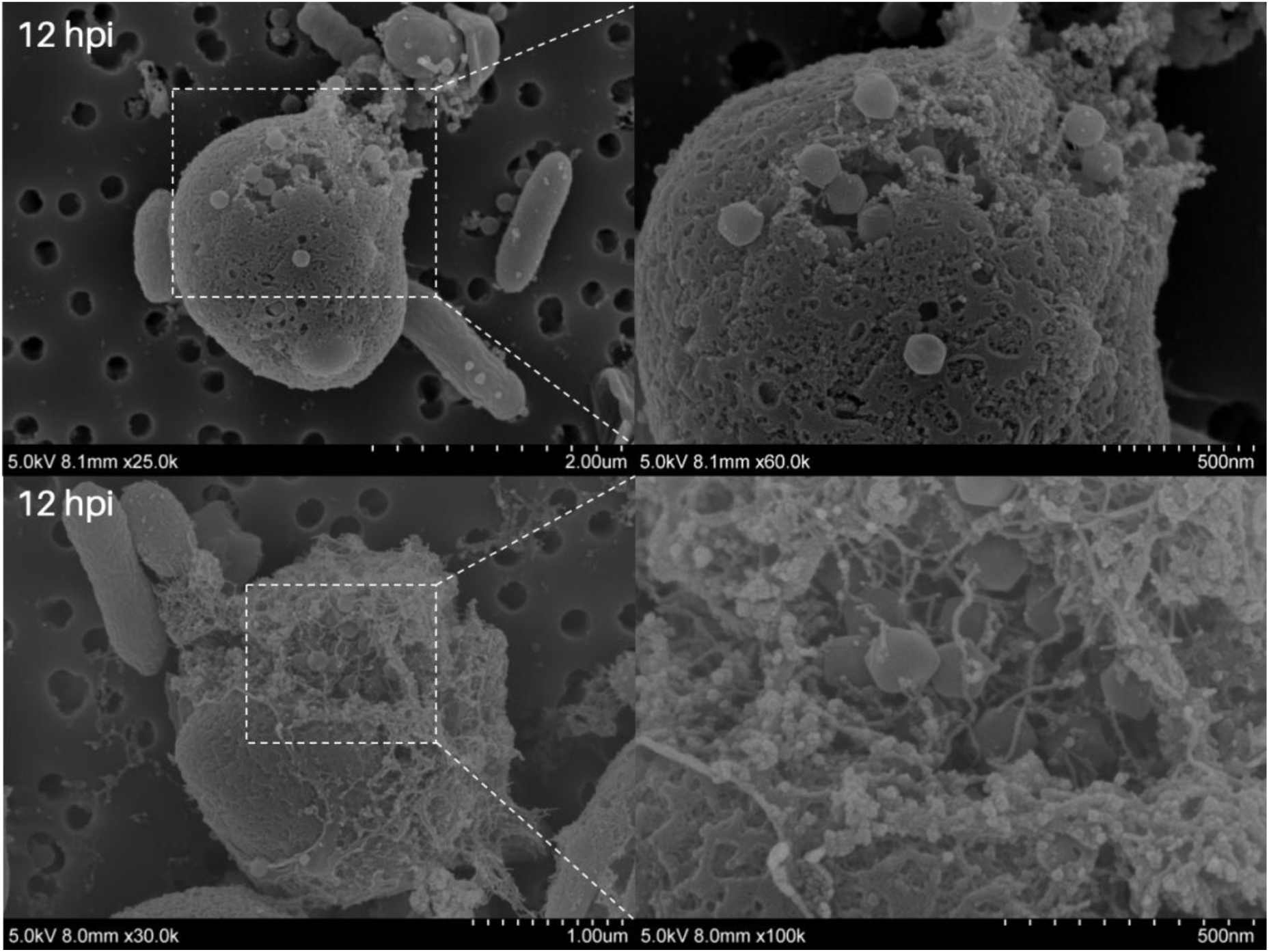
Non-tailed virions produced during egress. Scanning (SEM) electron micrographs of the *Pelagodinium*-PelV1 system at the late stage of infection

### Genomic features of PelV-1 and rare co-occurring virus, co-PelV

Our virus-host system is not axenic (contains bacteria and phages) so even after gradient purification, we process the sequences from targeted virion fractions as a metagenome to identify sequences from the dominant eukaryotic virus. Of the 16 bins produced by MetaBAT2, three could not be identified, eleven, were bacterial, and two were identified as viruses **(S1, S2 Table)**. The first viral bin, PelV-1, accounted for the majority of all reads (>80%) in the targeted iodixanol gradient fractions and the second, co-PelV, recruited very few reads (<1%) in those fractions. The dominance of PelV-1 over co-PelV sequences was also observed in recruitment of reads from unfractionated lysate (37% for PelV-1 and 0.0005% for co-PelV) **(S3 Table)**. Both viruses encode the ten commonly recognized NCLDV core genes, including MCP (major capsid protein), PolB (DNA polymerase B), A32 (A32-like packaging ATPase), VLTF3 (virus-like late transcription factor), SFII (superfamily II helicase), RNAPL, RPANS (RNA polymerase large and small subunits), mRNAc (mRNA capping enzyme), RNR (ribonucleotide reductase) and D5 (D5 primase/helicase) **(Fig. 6, S4 Table and S5 Table)**. The least pure gradient fraction analyzed was CsCl fraction 11 (1.385 g/L) in which only 38% of the reads mapped to PelV-1, while OptiPrep U5 (unfiltered lysate, fraction 5; 1.1685 g/L) showed the greatest enrichment, with 89% reads mapping to PelV-1. At the highest enrichment for co-PelV (OptiPrep fraction F7), only represents 0.6% of the total reads. OptiPrep was able to better separate the bacterial contaminants from the viruses than CsCl gradient based on both EM observations **(S1 Fig, S2 Fig)** and sequencing **(S1 Table)**.

**Figure 6.**
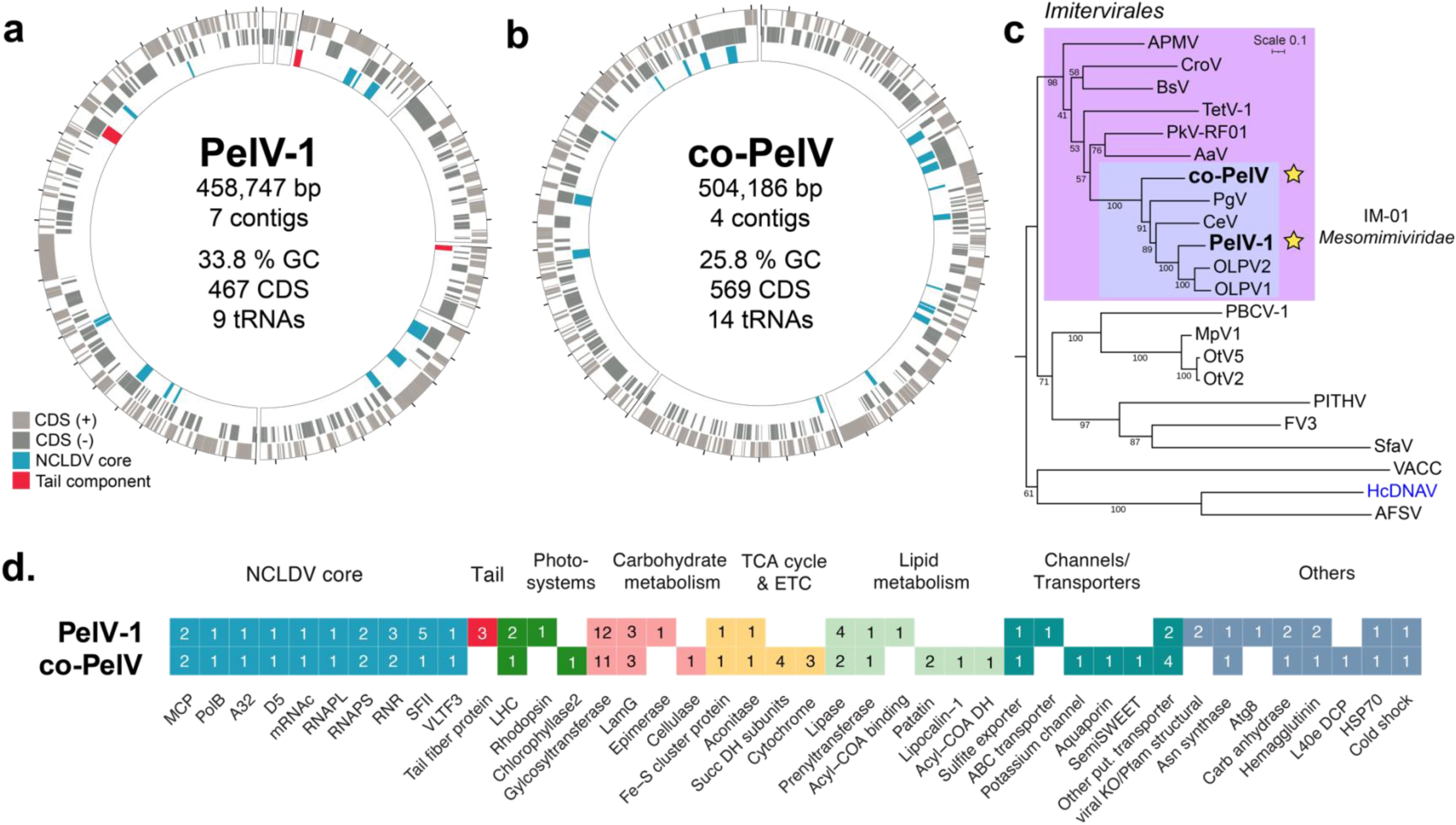
PelV-1 and co-PelV genome map and phylogeny. (a) PelV-1 and (b) co-PelV circularized genome map showing CDS/genes on positive and negative strands (gray boxes). NCLDV core genes (blue) and putative tail components (red) are labeled. (c) NCLDV phylogeny based on the PolB gene reveals that both viruses belong to Order Imitervirales, Family IM-01 (*Mesomimiviridae*). Phylogeny using other NCLDV markers is available in the supplementary. The first isolated dinoflagellate-infecting NCLDV, HcDNAV, is divergent (genus Dinodnavirus; in blue) and, as previously reported, clustered with the *Asfavirales* (ASFV). (d) Virologs and auxiliary metabolic genes are present in both genomes. Boxes show presence/absence and number of homologs.

In the PelV-1 genome (459 kb, 7 contigs, 33.8% GC) we identified 467 CDS/genes and 9 tRNAs, and in the co-PelV genome (504 kb, 4 contigs, 25.8% GC) we identified 569 CDS/genes and 14 tRNAs **(Fig. 6)**. Phylogenetic analysis using PolB and other markers shows PelV-1 and co-PelV belonging to order *Imitervirales*, family *Mesomimiviridae* (IM-01) **(Fig. 6C, S9 Fig).** We further implemented functional annotation against GVOG (Giant Virus Orthologous Groups), PFAM (protein families), KEGG orthologous groups, SwissProt, RefSeq, and NCBI nr (nonredundant proteins) databases, which show unique virologs (viral homologs) and various metabolic genes **(Fig. 6D, S4 Table, S5 Table).** About 23.7% (111 of 467) and 19.3% (110 of 569) of PelV-1 and co-PelV genes are not classified under a GV orthologous group, respectively **(S5 Table, S6 Table)**. Furthermore, 15.6% (73 of 467) and 5.27% (30 of 569) PelV-1 and co-PelV genes have no match to NCBI nonredundant protein database, respectively **(S4 Table & S5 Table)**. These sets of genes likely represent the ORFans, those without significant sequence similarity to known genes.

Like other NCLDVs, PelV-1 and co-PelV encode several copies of glycosyltransferases, methyltransferases, ubiquitin-related proteins, transporters, DNA replication and repair machinery, and transcription and translation factors. Other metabolic genes include those involved in amino acid, carbohydrate, and lipid metabolism, and components of TCA (tricarboxylic acid cycle), the electron transport chain, light harvesting/photosystem, and the cytoskeleton **(Fig. 6D)**. As specific examples, PelV-1 encodes for asparagine synthase (PelV1_244), sugar epimerase (PelV1_304), carbonic anhydrases (PelV1_384, 430), light-harvesting complex (PelV1_254), lipases (PelV1_314 & 387), viral rhodopsin (PelV1_132), aconitate hydratase (PelV1_358), ABC transporter (PelV1_120), iron-sulfur cluster accessory protein (PelV1_188), and autophagy-related protein Atg8 (PelV1_394).

The genome analysis shed light on putative tail components; PelV-1 encodes 3 genes that contain tail-like homologs, including a gene that matches *Synechococcus* phage tail fiber protein (PelV1_389), and two other tail-containing domain proteins (PelV1_011, 111) **(S10 Fig, S11 Fig)**. In contrast, the co-PelV genome does not encode tail-like protein homologs **(Fig. 6)**. Two of the PelV-1 genes are also potentially structural based on Pfam and KO annotations such as PelV1_042 (K25940: Poxviridae protein A33; Pfam: Xlink, hyaluronic binding), PelV1_052 (pfam02346: Vac_Fusion), and hemagglutinin homolog (PelV1_445) **(S4 Table)**. Phylogeny shows that PelV-1 rhodopsin belongs to the viral group 2 **(S12 Fig)**.

The co-PelV genome is equally interesting. Even though it does not encode for tail proteins, it contains putative surface proteins – hemagglutinin homologs (coPelV_180, 254). The co-PelV metabolic genes include carbonic anhydrase (coPelV_533), cellulase-like (coPelV_568), chlorophyllase (coPelV_079), cytochrome proteins C (co-PelV_149), cytochrome b6-f complex (coPelV_354) and ubiquinol-cytochrome C chaperone (coPelV_509), DMSP processing protein (coPelV_068), ion channel, (coPelV_161), light harvesting complexes (coPelV_277 & 426), lipases (coPelV_004, 121, 218 & 324), lipocalin (coPelV_468), and ribosomal L40e domain containing protein (coPelV_052). The co-PelV genome also encodes for key enzymes in TCA cycle such as aconitate hydratase (coPelV_182) and the complete succinate dehydrogenase subunits SDHA, SDHB, SDHC, and SDHD (coPelV_420, 423, 421, 422, respectively), as well as several transporters such as semiSWEET sugar transporter (coPelV_160) and aquaporin (coPelV_093). The PelV-1 and co-PelV LHC genes (with chlorophyll a/b binding domain) forms a cluster with other Mesomimivirids and SAR clade (**Fig. 7)**.

**Figure 7.**
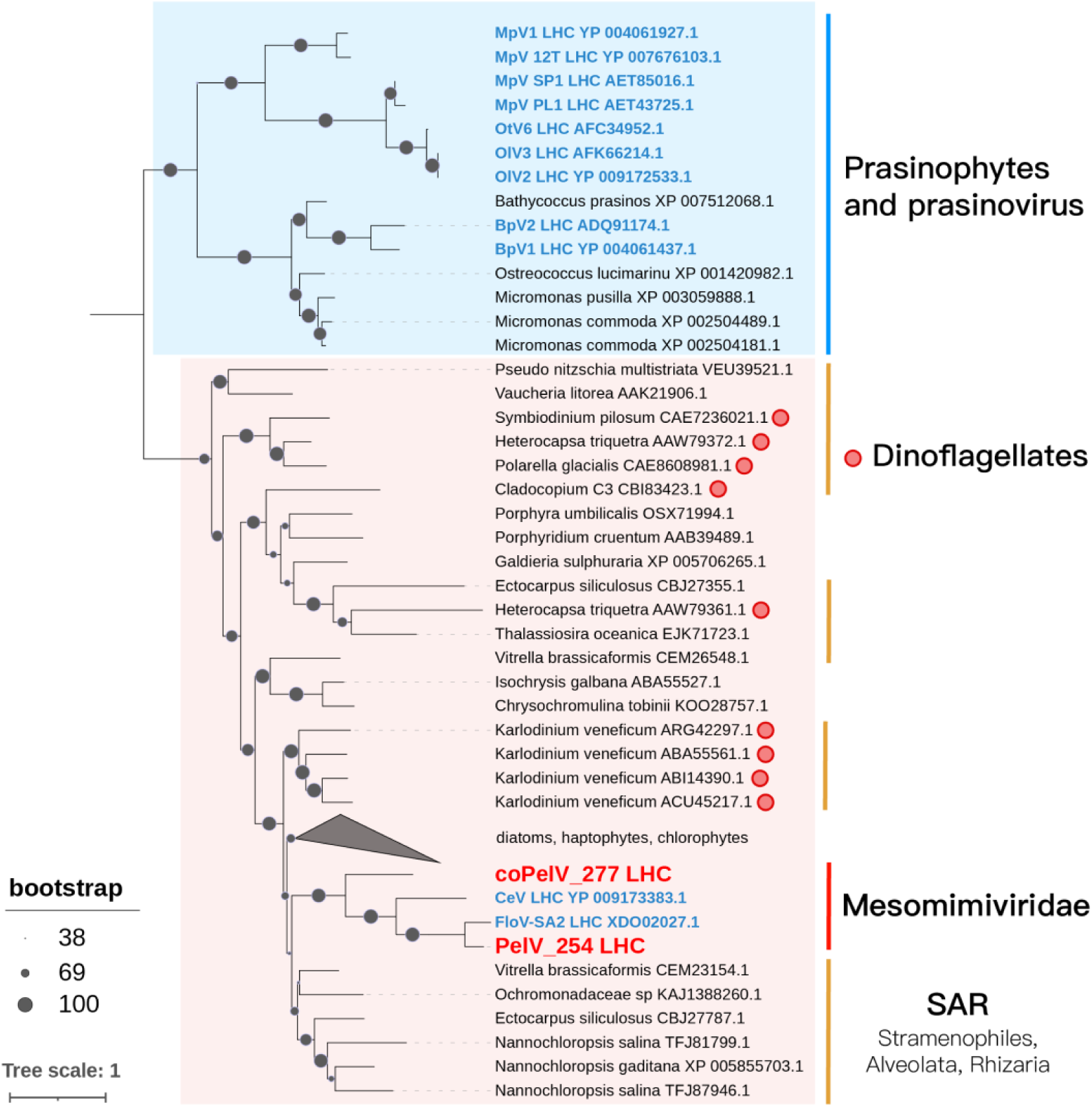
Phylogenies of PelV-1 and co-PelV chlorophyll AB domain-containing proteins (of light-harvesting complex; LHC). The tree includes proteins identified by Schulz et al. (2020) and additional BLASTP hits homologous to PelV-1 and co-PelV LHC sequences. Two main clusters are evident, first the prasinophytes and prasinovirus and second mainly SAR clade members and mesomimiviridae. Viral sequences are shown in blue, PelV-1 and co-PelV genes in red, and protistan sequences in black.

Analysis of the geographic extent of PelV-1 and co-PelV against global metagenomics data shows that both are widely distributed across the ocean at various depths (**Fig. 8)**. We showed the presence in both the surface and DCM layers but there are 3 occurrences in the mesopelagic layer as well. Most of the detections (47 of 59) were observed in the pico-size fraction (0.22 µm to 1.6 or 3 µm).

**Figure 8.**
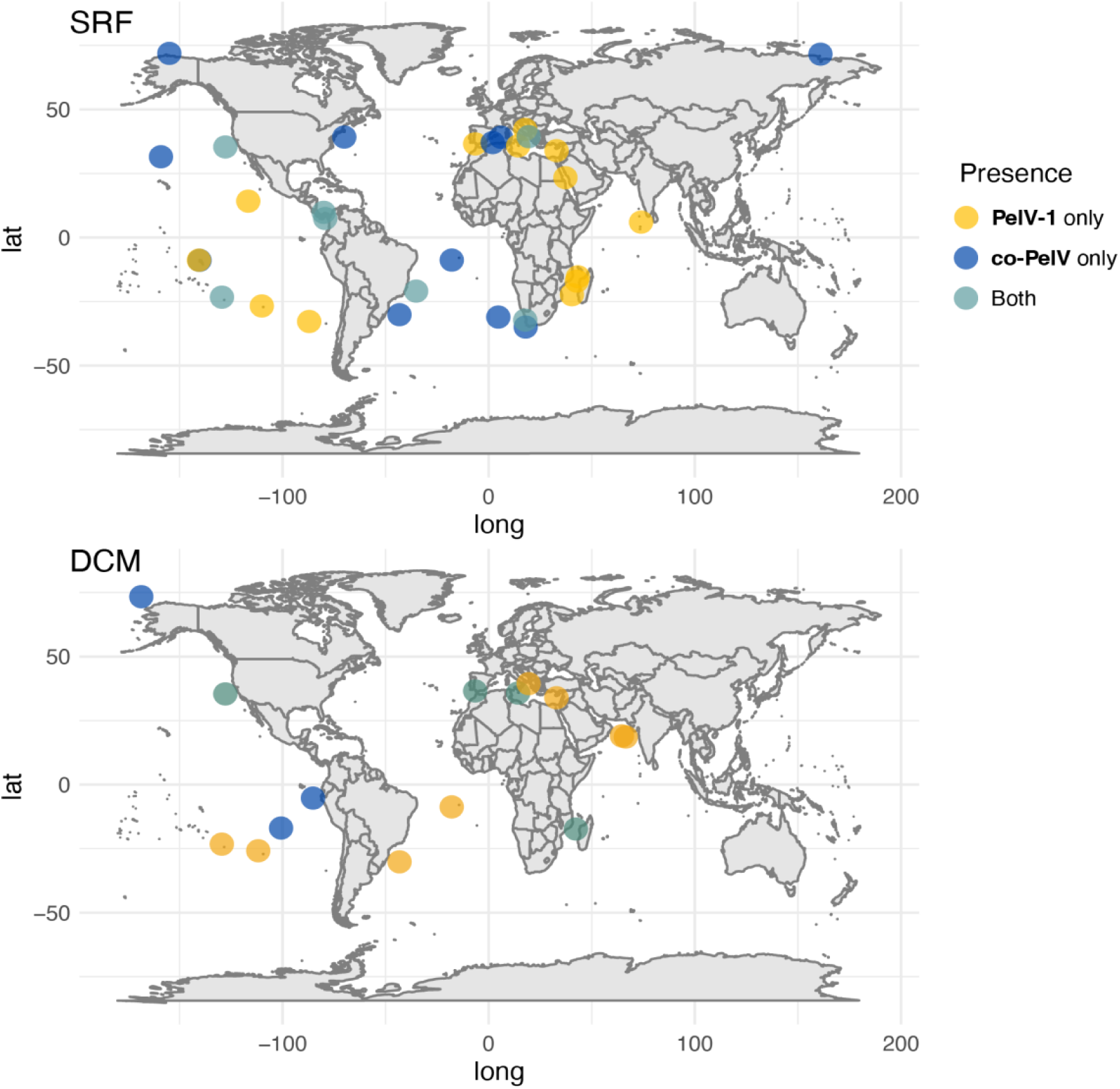
Global occurrences of PelV-1 and co-PelV phylotypes. PelV-1 and co-PelV phylotypes are detected across different oceanic regions and biomes.

### Quantifying PelV and co-PelV by qPCR

After finding evidence of a second genome present at apparent low abundance in the lysate, we prepared and fractionated a fresh lysate in an OptiPrep gradient. Quantification of the fractions by qPCR revealed several fractions with PelV-1 centered with a peak at a density of 1.163 g mL^-1^ **(S13 Fig).** The two fractions forming the bulk of the peak contained a total of 2 x 10^6^ copies. Negative staining of the two fractions with highest signal confirmed the presence of either the thin, but short (apparently broken), tail or possibly a stubby protrusion **(S13 Fig).** None of the fractions were positive for co-PelV either in the first round of qPCR (< 2 x 10^4^ copies) or a second round using more DNA to lower the detection threshold (< 2 x 10^3^ copies) suggesting that, if co-PelV was present in the second lysate, it made up < 0.1% of the virions. DNA extracted from uninfected *Pelagodinium* cells also did not test positive for either virus.

## Discussion

In this study, we report on the morphology and genome of a pleiomorphic micron-length tailed giant virus isolate, PelV-1, and the genome of a low abundance, co-occurring virus, co-PelV. The assembly of two distinct genomes from our culture was unexpected, but such co-culture is not unprecedented for giant viruses. Clandestinovirus, for example, was discovered in co-culture with Faustovirus ST1 grown on amoeba host *Vermamoeba vermiformis*, and was subsequently isolated (Cherif Louazani et al., 2017; Rolland et al., 2021). Although we had sufficient sequencing depth to assemble the co-PelV genome, this virus appears to have been present at very low levels compared to the dominant PelV-1. Based on relative numbers of reads from the whole lysate and various fractions sequenced, co-PelV abundance was ≤ 0.7% of PelV-1. In an attempt to separate the two viruses, a second lysate was fractionated, but this also supported the rarity of co-PelV (< 1 % of PelV-1 abundance). The failure to detect the virus in DNA extracted from an uninfected host suggests that co-PelV is not an endogenous virus that activates during infection of *Pelagodinium* with PelV-1. Because of the very low levels co-PelV, we could not isolate and further characterize the virus at this time, and the nature of its virion remains unknown. With PelV-1 so strongly dominating the lysate, we presume the various documented virion morphologies (all common) are just different forms of PelV-1. With co-PelV representing only one out of a hundred virions, it would be too scarce to establish as any of the documented common morphotypes. The variable morphologies we attribute to PelV-1 are presumably caused by triggered or spontaneous maturation processes post-lysis, or physical damage to some virions.

### Unique viral morphology and tail-like appendage

The most striking feature of the putative PelV-1 virion is the exceptionally long, thin tail-like structure observed on virions in fresh, unprocessed lysates. At its maximum (2.3 µm), this tail is longer than that seen on any other virus exceeding the ∼875 nm tail of the longest phage, P74-26 infecting *Thermus thermophilus* (Agnello et al., 2023) and another giant virus, Tupanvirus (tail at 0.55 - 1.85 µm) (Abrahão et al., 2018). Despite differences in absolute tail length, the ratio of tail length:capsid diameter ratio for PelV-1 (at the observed max length) and P74-26 phage is about the same, ∼11-12. Whether this is coincidental or whether some underlying mechanism constrains this ratio regardless of taxa is unknown. The observed considerable variations in tail length may result from tails being at different stages of development, from damage during purification, or both. The long, thin tails do appear to be easily broken as they are only commonly observed on virions in fresh lysate that has not been filtered or concentrated, similar to the observation in a tailed Ostreococcus virus, OtV09 (Thomy, 2022).

Although the tail of PelV-1 seems to be the longest described for a virus isolate, long-tailed virus-like particles have been reported on many occasions from different sources. Giant virus-like particles with tails >1 µm in length were observed in seawater from the North Atlantic (Bratbak et al., 1992) and North Pacific Oceans (Cochlan et al., 1993), in Antarctica sea ice (Gowing et al., 2002), and soil samples (Fischer et al., 2023). Tailed virus-like particles were also observed inside several green algal cells [reviewed in (Van Etten et al., 1991)], and in cultures of the filamentous green alga, *Uronema gigas* (Dodds & Cole, 1980). We hypothesize that the long tail appendage of PelV-1 confers an advantage as it increases the effective diameter of the virus, increasing the probability of contacting a potential host.

Increased contact rates could prove particularly beneficial in low biomass systems such as the oligotrophic North Pacific Ocean. *Micromonas pusilla* Naples virus 1 and 2 (MpNV1, MpNV2), which also infect small flagellate phytoplankton, have been shown to contain a short tail used for host attachment (Zingone et al., 2006). Compared to eukaryote-infecting viruses, more is known about the function of tails in bacteria-infecting viruses. For example, phage tails are crucial in host surface recognition, penetration and delivery of genetic material [reviewed in (Nobrega, 2018)]. It has also been shown that tail fibers can fold back upon attachment (B. Hu et al., 2015). The host-attached virions observed by electron microscopy are attached by a short tail, but whether this resulted from contraction or depolymerization of an initially long tail, or the attachment is by the stubby appendage is unknown.

Another prominent morphological feature of PelV-1 is the “stargate” apex opening, a common attribute of mimiviruses such as the type species Acanthamoeba polyphaga mimivirus (Mutsafi et al., 2014; Zauberman et al., 2008), Bodo saltans virus (Deeg et al., 2018), *Cotonvirus japonicus* (Takahashi et al., 2021), and Tupanvirus (Abrahão et al., 2018). Likewise, the nucleoprotein-like assembly found near the PelV-1 capsid is similar to that of Acanthamoeba polyphaga mimivirus (ApMV) (Villalta et al., 2022). However, we cannot rule out that these nucleoprotein-like assemblies could also originate from the dinoflagellate host, as dinoflagellates are known to possess condensed genomes exhibiting similar morphologies (Gornik et al., 2019; Soyer-Gobillard et al., 1996).

During an infection time course, we observed that PelV-1 uses its tail for attachment at 1 mpi to 6 hpi (minute- or hour post-infection) and the entry mechanism appears to be endocytosis-like, potentially phagocytosis, consistent with previous results showing that this strain of *Pelagodinium* is capable of phagotrophy (Q. Li et al., 2022). Lysed cells at 12 hpi showed the production of a non-tailed morphotype. Numerous ultrathin sections from two separate time-series experiments also confirm the absence of a tailed morphotype inside the cell during the late stage of infection. This is similar to the report of the tailed OtV09, wherein the non-tailed variant was observed inside *Ostreococcus* cells across an infection time course (Thomy, 2022). This suggests that the tail develops outside the host. Such post-lysis morphogenesis is reminiscent of that observed for the *Aciadianus* two-tailed virus (Häring et al., 2005). This could also explain the varying tail length seen in lysates. A bacteriophage in the family *Autolykiviridae* was shown to develop a tail upon contact with the host cell (Kauffman et al., 2018). Self-assembling viral proteins are common and include capsids (Buzón et al., 2020; Chaudhary & Yadav, 2019) and giant virus histones (Irwin & Richards, 2024). Although host-independent tail assembly is plausible, this still requires experimental evidence to confirm.

### Genome and metabolic capacity of PelV-1 and co-PelV

Viral purification and sequencing revealed PelV-1 and co-PelV genomes with unique metabolic capacities. PelV-1 and co-PelV are comparable in genome size and gene content to other mesomimivirus, such as Chrysochromulina ericina virus (CeV, 460kb, 434 genes, 32% GC) and Phaeocystis globosa virus (PgV, 474kb, 512 genes, 26.3% GC) (Gallot-Lavallée et al., 2017; Santini et al., 2013). TCA cycle genes have been found in the metagenomics samples (Moniruzzaman et al., 2020) as well as in isolates, PkV-RF01 (Blanc-Mathieu et al., 2021) and *Pandoravirus massiliensis* (Aherfi et al., 2022). Here we show that both PelV-1 and co-PelV contain several key enzymes for the TCA cycle such as aconitate hydratase. Similar to PkV-RF01, co-PelV contains all four subunits of succinate dehydrogenase and three cytochrome proteins. This suggests the use of TCA cycle genes to meet the energy demand of infection, potentially as alternative when the host shuts down transcription or translation (Blanc-Mathieu et al., 2021) or as viral genes replace suppressed host genes (Derelle et al., 2018).

Both PelV-1 and co-PelV encode for a cold-shock protein, a known RNA chaperone that prevents secondary structure formation and misfolding (Keto-Timonen et al., 2016; Phadtare et al., 1999; Rudan et al., 2015), and HSP70 (heat shock protein 70), central to protein homeostasis (C. Hu et al., 2022). Both stress-related genes might confer some level of protection to the virocell during host-induced stress response/autophagy. Another autophagy-related gene encoded by PelV-1, Atg8, is a key component in the formation of autophagosomes (Shpilka et al., 2011). Whether it performs autophagy or non-canonical functions, such as cargo degradation and secretion (Nieto-Torres et al., 2021), remains to be seen.

Phylogenies of PelV-1 and co-PelV light harvesting complex (LHC) gene indicates possible acquisition from a member of the SAR clade within the eukaryotes, similar to the situation for CeV (Gallot-Lavallée et al., 2017). The presence of host-derived photosynthesis genes in eukaryotic viruses is reminiscent of the widespread prevalence of host-derived cyanophage photosystems (Lindell et al., 2005; Luo et al., 2020; Sullivan et al., 2006), illustrating a common theme of adaptation across diverse groups of viruses. LHC proteins, specifically the chlorophyll a/b binding protein, were shown to play a role in high-light and oxidative stress (Staneloni et al., 2008; Yang et al., 1998). Another light-harvesting protein in PelV-1 is rhodopsin, which belongs to viral rhodopsin group II (VRGII). Functional characterization of VRGII has shown they act as a light-gated pentameric ion channel (Bratanov et al., 2019), and the PelV-1 rhodopsin may function similarly. The potassium channel encoded by co-PelV could function similarly to the PBCV-1 potassium channel, which triggers depolarization, allowing DNA injection into the host cell (Lee et al., 2016). A similar channel encoded by another marine algal virus isolate, TetV-1 (Schvarcz & Steward, 2018), was found to be functional (Kukovetz et al., 2020).

PelV-1 and co-PelV both possess asparagine synthetase, which is widespread in mimiviruses (Mozar & Claverie, 2014). The function of this gene may be associated with a proteome bias for amino acids that include asparagine residues, as shown in Acanthamoeba polyphaga mimivirus (Raoult et al., 2004). PelV-1 and co-PelV also both possess various glycosyltransferases, which are widely observed in NCLDVs, and are likely utilized in viral protein glycosylation, as observed in mycobacteriophages (Freeman et al., 2023) and PBCV-1 (Graves et al., 2001).

A number of other notable genes are encoded by co-PelV. To our knowledge, chlorophyllase2, though detected in MAGs (Schulz et al., 2020), has not yet been described in a virus isolate. Cellulase-like glycosyl hydrolase and aquaporin are also potentially novel genes not yet seen in cultivated viruses. Chlorophyllase2 might be involved in breaking down chlorophyll and may provide energy or material for virions, but at the expense of absorbing light. Cellulase can potentially play a role in the degradation of host cell wall. On the other hand, aquaporin, a water channel, is implicated in the cellular inflation of the dinoflagellate *Pyrocystis noctiluca*, allowing the remarkable vertical migration in the North Pacific Subtropical Gyre (Larson et al., 2022). This suggests that giant viruses can potentially influence not only the host metabolism but also their behavior. Whether the extremely low abundance co-occurring co-PelV can be cultured separately remains to be seen and a subject of a future effort.

In conclusion, here we presented the annotated genomes of two new members of the family *Mesomimiviridae,* PelV-1 and co-PelV, and describe the unusual morphological traits of the former, PelV-1, which has the longest tail of a cultivated virus described to date. The genome annotations illustrate the immense metabolic capacity of viruses, with genes involved in carbohydrate metabolism, TCA cycle, cytochrome, photosynthesis, and rhodopsin, among others. The systems described here are a promising emerging model system to investigate dinoflagellate-virus dynamics and the mechanism and consequences of tail formation in a eukaryote-infecting virus.

## Materials and Methods

### Host isolation and identification

The *Pelagodinium* sp. isolate (UHM4000) was cultivated from seawater collected at 25 meters depth from Station ALOHA (22°45’ N, 158°00’ W), 100 km north of Oahu, Pacific Ocean on May 4, 2019 during Cruise #311 of the Hawaii Ocean Time-series program (Karl & Church, 2014; Karl & Lukas, 1996). The strain was rendered unialgal, but not axenic, through repeated serial dilution-to-extinction. The strain can be maintained autotrophically (K medium with no added prey) or mixotrophically (K minus N medium, amended with *Prochlorococcus*) (Q. Li et al., 2022). The strain was identified by gross morphology and 18S rRNA gene sequence (GenBank ID# MZ611719). The culture was deposited at the National Center for Marine Algae and Microbiota (NCMA) as previously described (Q. Li et al., 2022), but, as of this publication date, has not been assigned an NCMA collection number. We used the autotrophically grown host for the rest of the experiments to obtain a dense culture amenable to counting and molecular analyses.

### Virus isolation

The addition of unfiltered seawater (200 mL) collected at 25 m at Station ALOHA on July 17, 2020, to a *Pelagodinium* host culture (1 L) resulted in cell lysis. The lytic agent was then subjected to three rounds of dilution-to-extinction in 96-well plates, as previously described (Nagasaki & Bratbak, 2010). The virus was maintained by monthly transfer of lysate to fresh host culture.

### Virus purification using CsCl density gradient

A viral lysate (950 mL) was made by adding 10-20% by volume of previously produced lysate to healthy *Pelagodinium* sp. culture in March 2022. After 7 days, the lysate was passed through a 0.65 µm pore-size, polyvinylidene difluoride (PVDF) membrane filter to remove cell debris and some bacteria. The virus was then concentrated by tangential-flow filtration (30 kDa TFF; Pellicon XL PXC030C50). To enhance recovery, a backflush-with-recirculation step was added at the end of the concentration phase. To do this, a 10-cc syringe filled with permeate was attached to the permeate line near the end of the concentration step, being careful to avoid bubbles in the line. While recirculation continued, pressure was applied to the syringe plunger to provide a slow reverse flow (2–5 mL min^-1^) across the membrane. After recovery, from the ultrafilter, the concentrate was filtered a second time (0.65 µm PVDF) and further concentrated using Amicon Ultra-15 (100 kDa). The virus was purified on a continuous equilibrium buoyant density gradient prepared with CsCl as previously described (J. E. Lawrence & Steward, 2010) **(S1 Fig)**. Briefly, an initial step gradient of 1.20, 1.45, 1.60 g/L (SM buffer as diluent), with sample incorporated into the middle density fraction, was centrifuged in an ultracentrifuge (SW 41 Ti rotor; 45 hrs; 30,000 RPM, 4^°^C). After centrifugation, positions of visible bands were recorded, and gradient fractions (ca. 500 µl) were recovered using LabConco Auto DensiFlow. The density of each fraction was determined by the change in fraction weight after removing a known volume with a positive displacement pipet (J. E. Lawrence & Steward, 2010). Fractions were checked for the presence of viruses by transmission electron microscopy (TEM) as described below.

### Virus purification for metagenomics and microscopy in an iodixanol density gradient

We performed additional density gradient fractionations, this time with iodixanol (OptiPrep^TM^, Serumwerk Bernburg AG), to obtain purer fractions. Four liters of lysate was generated by adding 10% by volume of previously produced lysate to healthy *Pelagodinium* sp. culture in November 2023. After 2 days, a portion of the lysate (3 L) was pressure filtered via peristaltic pump through a stack of a glass-fiber filter (125 mm GF/C with nominal pore size of 1.2 µm; Cytiva) overlaid on a polyvinylidene difluoride membrane filter (PVDF, 142 mm, 0.45 µm Durapore; Millipore-Sigma) inside a stainless steel 142 mm filter holder. Viruses in the filtered lysate were concentrated into a smaller volume via tangential flow filtration (30 kDa TFF; Pellicon XL PXC030C50). We further concentrated the viruses using a 50 kDa Amicon Ultra-15. For remaining unfiltered lysate (1 L), the viruses were concentrated into smaller volume using centrifugal ultrafiltration devices (100 kDa Centricon Ultra; Millipore Sigma). Pre-formed OptiPrep continuous gradients were prepared with a two-chambered gradient maker connected to the Auto DensiFlow with low-(20%) and high-density (45%) end member solutions prepared in 0.2 µm filtered autoclaved filtered seawater. Concentrated lysates were layered on top of the pre-formed gradients and centrifuged at 35,000 RPM (210,000 x g; SW 41 Ti rotor) at 20°C for 21 hrs. The collection of fractions, checking of their density, and screening for presence of viruses were as described for the CsCl gradient **(S2 Fig.)**. One difference was that fraction volume was allowed to vary in an effort to ensure that each visible band was collected in its own fraction and that all of any given visible band was in a single fraction.

### DNA extraction and sequencing of purified viruses

Fractions from CsCl (n = 2 fractions) and OptiPrep gradient (n = 4 fractions) that were confirmed by TEM to have high virus concentration and limited bacterial contamination were prepared for extraction by buffer exchange with SM buffer (for CsCl) and seawater (OptiPrep) in centrifugal ultrafilters (Microcon, Millipore Sigma). DNA was then extracted by a spin-column method (DNA MicroPrep kit, Zymo) as described in the manufacturer’s protocol. We added a proteinase K lysis step for fractions obtained from the OptiPrep gradient. DNA concentration was measured using Qubit. Then, samples were sent to SeqCenter, Pittsburgh, PA, for 2 x 151 bp PE Illumina sequencing. We sequenced at 200 Mb- and 1 Gb-scale for CsCl and OptiPrep fractions, respectively.

### Genome analysis

Raw sequences were trimmed and quality controlled using Trimmomatic (with following criteria Leading:30 Trailing:30 SlidingWindow:4:15 MinLen:30) (Bolger et al., 2014). All sequences were concatenated prior to MEGAHIT assembly (D. Li et al., 2015). We then performed Bowtie2 mapping (Langmead & Salzberg, 2012) and MetaBAT2 binning (Kang et al., 2019). DAS tool was used to identify the contigs-to-bin memberships (Sieber et al., 2018), and SeqKit was used to determine the length statistics (Shen et al., 2016). Our focus was on identifying Nucleocytoplasmic Large DNA Viruses (NCLDVs), the group of viruses comprising the phylum *Nucleocytoviricota*. Initial identification of *Nucleocytoviricota* (or NCLDV for short) contigs was done using geNomad (Camargo et al., 2023) and the resulting bins confirmed using ViralRecall (Aylward & Moniruzzaman, 2021). We identified an overwhelmingly dominant NCLDV, which we called PelV-1 (up to 89 % of total sequencing reads of the purified viral fraction), and a rare co-occurring NCLDV, which we named co-PelV (<0.8 % relative abundance compared to PelV-1) **(S1, S2, S3 Table).** The relative abundance of bins was calculated by dividing the number of recruited reads by the number of raw sequencing reads. The taxonomy of the other non-viral bins was estimated in Anvi’o using the anvi-estimate-scg-taxonomy function (Eren et al., 2015) based on the Genome Taxonomy Database (Parks et al., 2018, 2020). Gene identification was done using Prodigal (-p meta) (Hyatt et al., 2010), while initial functional annotation was done using ViralRecall using PFAM and GVOG databases (Aylward & Moniruzzaman, 2021). Further functional annotation was implemented in MicrobeAnnotator using KEGG orthology (release 109.1), SwissProt (2024_01 release), RefSeq (release 223), and TrEMBL (2024_01 release) databases (Ruiz-Perez et al., 2021). We also performed Diamond blastp against NCBI nonredundant (nr) protein database (Altschul et al., 1990; Buchfink et al., 2021; Sayers et al., 2022). MicrobeAnnotator KO annotations were found to be lenient, so for confirmation, KO annotations were implemented using the eggNOG (http://eggnog-mapper.embl.de/) (Huerta-Cepas et al., 2019) and KofamKOALA webservers (https://www.genome.jp/tools/kofamkoala/) (Aramaki et al., 2020). Finally, BioCircos in R was used for the genome visualization (Cui et al., 2016)

### Virus fractionation in an iodixanol density gradient to confirm PelV-1 morphology

After detecting a second viral genome in the initial lysates, a new lysate was generated, fractionated, and analyzed by qPCR and electron microscopy to determine the relative abundance and buoyant densities of the dominant and minor contaminating virus. Seventy-five ml of lysate was generated by adding 10% by volume of previously produced lysate to healthy *Pelagodinium* sp. culture in April-May 2024. The lysate was passed via a peristaltic pump through a GF/C filter overlaid on a 0.65 µm PVDF membrane filter. Viruses in the filtered lysate were concentrated into a smaller volume via centrifugal ultrafiltration (30 kDa Centricon-Plus-70 mL; Millipore Sigma). Pre-formed OptiPrep gradients were prepared as described above. Concentrated filtered lysate was layered on top of the pre-formed gradient and centrifuged at 35,000 RPM (210,000 x g; SW 41 Ti rotor) at 20°C for 22 hrs. Fractions (400 µL each; 38 total fractions) were collected with a piston fractionator (Biocomp Instruments). Gradient medium was removed from the fractions and the volume reduced by washing with filtered seawater in centrifugal ultrafilters (30 kDa Microcon; Millipore Sigma). A portion (10 µL) of each concentrated, seawater-exchanged fraction was fixed (0.5% glutaraldehyde) for electron microscopy, and the remainder saved for DNA extraction.

### Quantifying PelV-1 and co-PelV by quantitative PCR (qPCR)

DNA was extracted from half (250 µL) of the seawater-exchanged fractions using a spin-column method (Zymo Research Quick-DNA/RNA Viral Kit) and from *Pelagodinium* cells pelleted from 10 mL of an uninfected culture (Zymo Research Quick-DNA Microprep Kit). DNA concentration in the extracts was measured fluorometrically using Qubit (Thermo Fisher). Primers targeting the DNA polymerase gene (PolB) and specific to either PelV-1 or co-PelV were designed via manual inspection of the aligned sequences in Geneious to find areas of difference a suitable distance apart (150 to 300 bp). Candidate primer sets were screened *in silico* with the OligoAnalyzer tool on the Integrated DNA Technologies (IDT) website to identify those with a low tendency to form hairpins, self-dimers, or hetero-dimers. The selected primers range from 20–23 nt **(S6 Table)** and each pair is expected to produce an amplicon around 200 bp in length (194 bp for PelV-1, 209 bp for co-PelV) in the same region of the PolB gene of each virus (**S14 Fig**). The primers have sufficient mismatches to the corresponding non-target virus (7 to 11, including mismatched 3’ terminal bases) to ensure specificity. A synthetic standard (500 nt gBlock™) for qPCR analysis of either target was designed by including forward and reverse primer target sites for each virus (**S15 Fig**) arranged so that each pair produces an amplicon similar in size (201 or 205 bp for PelV-1 and co-PelV, respectively) to that of the natural target. Aside from the primer sites, the rest of the gBlock sequence (upstream, downstream, and in between the primer sites) was designed by initially copying from the corresponding locations of the consensus PolB sequence (randomly choosing between the residues in PelV-1 or co-PelV where they differed), then further altering the sequence manually to eliminate some homopolymer stretches to meet the IDT recommendations for complexity. A suitable annealing temperature for the primers was first empirically tested by standard PCR using the same reagents that would be used for qPCR (below) and a gradient block thermal cycler (BioRad iCycler).

Subsequent qPCR reactions consisted of 10 µl of PowerTrack SYBR Green Master Mix, 1 µl each of 300 nM forward and reverse primers, 5 µl of nuclease-free water, and 3 µl of DNA. Cycling was done in a QuantStudio 3 thermal cycler with real-time monitoring (Thermo Fisher). Two biological and two technical replicates were assayed along with a negative (non-template) control and a standard curve consisting of a dilution series of the gBlock (6×10^7^–6×10^2^ copies). PCR cycling conditions were 95°C for 3 minutes followed by 30 cycles of 95°C for 30 seconds, 56°C for 30 seconds, and 72°C for 30 seconds with a continuous melt curve at 60°C-95°C with 1.6°C/s increments. Primer efficiency and primer specificity were assessed from standard curves **(S16 Fig)** and melt curves, respectively. To improve the detection limit, the assay was run a second time after concentrating the gradient fraction extracts 2.5-fold using centrifugal ultrafiltration (30 kDa Microcon; Millipore Sigma) and using 2.7 times more of the concentrated extract volume, resulting in a detection limit 6.7 times lower than the first round. The qPCR reactions in this case consisted of 10 µl of PowerTrack SYBR Green Master Mix, 1 µl each of 300 nM forward and reverse primers, and 8 µl of DNA. All other aspects of the qPCR were unchanged.

### Phylogenetic analysis

A PolB phylogeny was created by protein alignment using MAFFT (--localpair --maxiterate 1000), trimmed with trimal (-gappyout), and maximum-likelihood phylogenetic trees inferred using IQ-tree2, with automatic model selection, ultrafast bootstrap calculation, and SH-aLRT test and (-B 1000 -m MFP - alrt 1000). For PolB, LG+F+I+G4 was chosen as the best-fit model. PolB was used to include the known dinoflagellate NCLDV HcDNAV, for which only this marker gene was available. To confirm the placement of PelV-1 and co-PelV, additional phylogenetic trees were made using A32, Topo2, RNAPL, and VLTF3 markers extracted using NCLDV MarkerSearch (Moniruzzaman et al., 2020). Phylogenetic trees of rhodopsin and light-harvesting complex (LHC) were constructed using the same methods, except for rhodopsin tree which was inferred through Neighbor Joining. A wide variety of protistan, bacterial and viral rhodopsin, and LHC protein sequences were obtained from previous report (Schulz et al., 2020) and NCBI blastp matches. Trees were visualized in iTOL (Letunic & Bork, 2021). Protein alignment files are provided as supplementary materials.

### Presence of PelV-1 and co-PelV in global metagenomes

To assess the global distribution of PelV-1- and co-PelV-like viruses, we performed a local blastP of PolB sequences against the Tara Ocean NCLDV PolB phylotypes, as previously described (Endo et al., 2020). The top hits for PelV-1 (TARA_030_DCM_0.22-1.6_G_scaffold270213_1_gene279451; 91% identical, eval < 0.0005) and co-PelV (TARA_067_SRF_0.22-0.45_G_scaffold177636_2_gene190099; 88% identical, eval < 0.0005) were then matched to the occurrences of these phylotypes to the Tara Ocean metagenomes.

### Infection time series

To observe details of virion attachment, changes in virocell ultrastructure, and virion egress, subsamples were collected and preserved for electron microscopy over the course of an infection cycle. To determine which samples to prepare for electron microscopy, the progress of infection was monitored by tracking host abundance with flow cytometry (AttuneTM NxT; Thermo Fisher Scientific). The host was readily identified using side scatter (SSC) and chlorophyll autofluorescence, but a cytometric signature for PelV-1 could not be reliably distinguished from bacterial contaminants and cell debris. A preliminary time series was done in March 2022 to examine PelV-1 inside host cells. Two healthy and two infected cultures were monitored until 150 hours post-infection (hpi). Infected cultures were made by adding 100 mL fresh lysates to 550 mL exponentially growing *Pelagodinium* culture. At fixed intervals until 150 hpi, 1 mL sample was fixed using glutaraldehyde (1% v:v final conc.), flash frozen, and stored at -80°C until analysis. 50 mL samples, one for control and one at 12 hpi were fixed (final concentration: 0.1 M sodium cacodylate, 2% glutaraldehyde, 0.005 M CaCl_2_, 0.06 g glucose/mL) for agarose embedding, ultrathin sectioning, and TEM. A more detailed time series was carried out in November 2022. This time, triplicate control and triplicate infected cultures were monitored until 48 hpi. For the infected replicates, 200 mL of freshly produced lysate was added to 1 L of exponentially growing culture. For the controls, 200 mL of K-media was added to 1 L of culture. Samples for EM (50 mL) and flow cytometry (1 mL) were collected and fixed, 1 hour before infection, 15 minutes post-infection (mpi), 3 hpi, 6 hpi, 9 hpi and 12 hpi. Fixed samples for EM were pelleted at 2,400 x g for 10 minutes at 4°C. 1 mL was saved for ultrathin sectioning and transmission electron microscopy (see below), and another 1 mL was saved for scanning electron microscopy (see below).

### Electron microscopy of virions

Viral morphology was observed using negative staining. Briefly, 4 µl of the sample (raw viral lysate or purified density gradient fraction) was applied to a freshly glow-discharged Formvar-coated grid. Samples were carefully wiped with filter paper and washed with 10 µl distilled water. Four microliters of 2% uranyl acetate were applied for staining. Grids were then washed with 10 µl distilled water, dried before viewing on a Hitachi HT7700 TEM at 100 kV and photographed with an AMT XR-41B 2k x 2k CCD camera.

### Electron microscopy of thin sections

Fixed samples were mixed with 5% low-melt agarose at a 2:3 volume ratio. The agarose was then solidified on ice and cut into <1 mm blocks. The agarose-embedded samples were then fixed with 0.1 M sodium cacodylate buffer with 0.44 M sucrose twice for 20 minutes and followed by post-fixation with OsO_4_ in 0.1 M sodium cacodylate buffer for 1 hour. Samples were then dehydrated in a graded ethanol series (30%, 50%, 70%, 85%, 95%, 100%), substituted with propylene oxide, and finally embedded into LX112 epoxy resin. Ultrathin sections (60-80nm) were obtained on an RMC MT6000-XL Ultramicrotome, viewed unstained on a Hitachi HT7700 TEM at 100 kV, and photographed with an AMT XR-41B 2k x 2k CCD camera.

### Scanning electron microscopy

Samples in fixative were loaded on a 0.2 µm polycarbonate filter held in Swinnex filter holders and washed twice with 0.1 M sodium cacodylate buffer with 0.44 M sucrose, then postfixed with 1% OsO_4_ in 0.1 M sodium cacodylate, dehydrated through an ethanol series (similar to above), and dried in a Tousimis Samdri-795 critical point dryer. The filters were then mounted on aluminum stubs and sputter coated with gold/palladium in a Hummer 6.2 sputter coater. Samples were viewed using a Hitachi S-4800 Field Emission Scanning Electron Microscope with an accelerating voltage of 5kV.

## Acknowledgments

This project was funded by US National Science Foundation grants awarded to KFE and GFS (OIA #1736030 and OCE #2129697). The bioinformatics work in this paper was implemented at the University of Hawai‘i High-Performance Computer. The technical support and advanced computing resources from the UH Information Technology Services – Cyberinfrastructure, funded partly by the NSF MRI award # 1920304. This article reports data obtained at the University of Hawaiʻi at Mānoa Microscopy Core, supported by NIH NIGMS grants P20GM139753 and P20GM125508. We thank Victor Limon and Samantha Lopez for assisting in one of the time series samplings. The Uehiro Center for the Advancement of Oceanography Graduate Student Fellowship supports APG and ABL.

## Data Accessibility

The sequencing data used in this paper have been deposited in the NCBI Short Read Archive as BioProject PRJNA1256882.

## Author Contributions

APG, CRS, KFE, and GFS conceived of the study. APG, CRS, and ABL performed experiments and analysis. TMW and AIC performed experiments and contributed data. APG, KFE, and GFS wrote the manuscript. All contributed to editing and approving the final manuscript.

## Conflict of Interest

The authors declare no conflict of interest.

## Notes

### Competing Interest Statement

The authors have declared no competing interest.

