## Supplemental Material 1 for "A dinoflagellate-infecting giant virus with a micron-length tail"

### Supplementary Materials

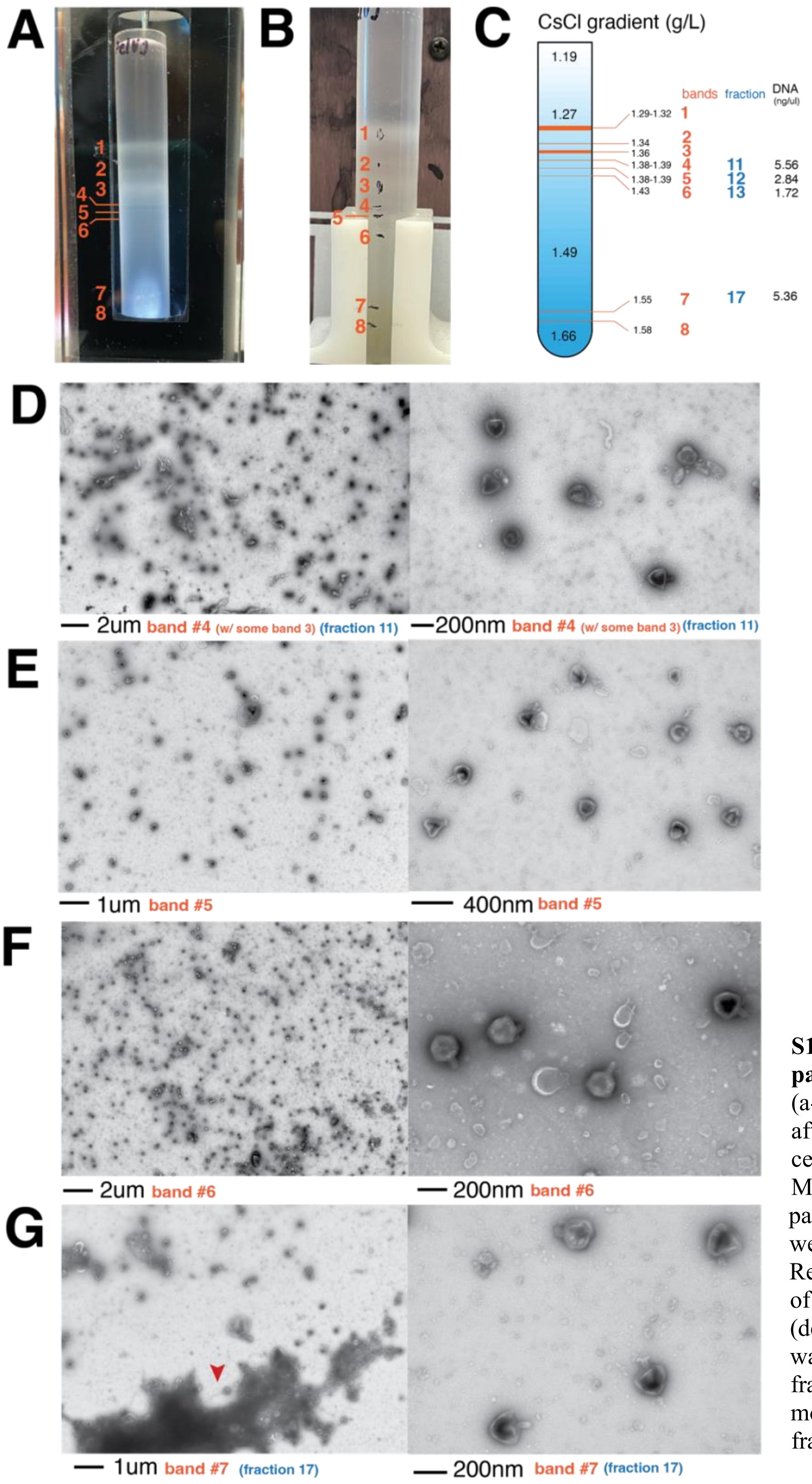


**S1 Figure. Purifying PelV-1 particles using a CsCl gradient.** (a-c) Several bands were observed after ultracentrifugation using a cesium chloride density gradient. Most of the concentrated PelV-1 particles, with rare contaminants, were observed in bands 3 to 6. (d-g) Representative electron micrograph of purified PelV-1 from bands 3-6 (density of 1.36 – 1.43 g/L. DNA was extracted and sequenced from fractions 11 and 17. (g) Long-tail morphotype was also observed in fraction 17 (red arrow).

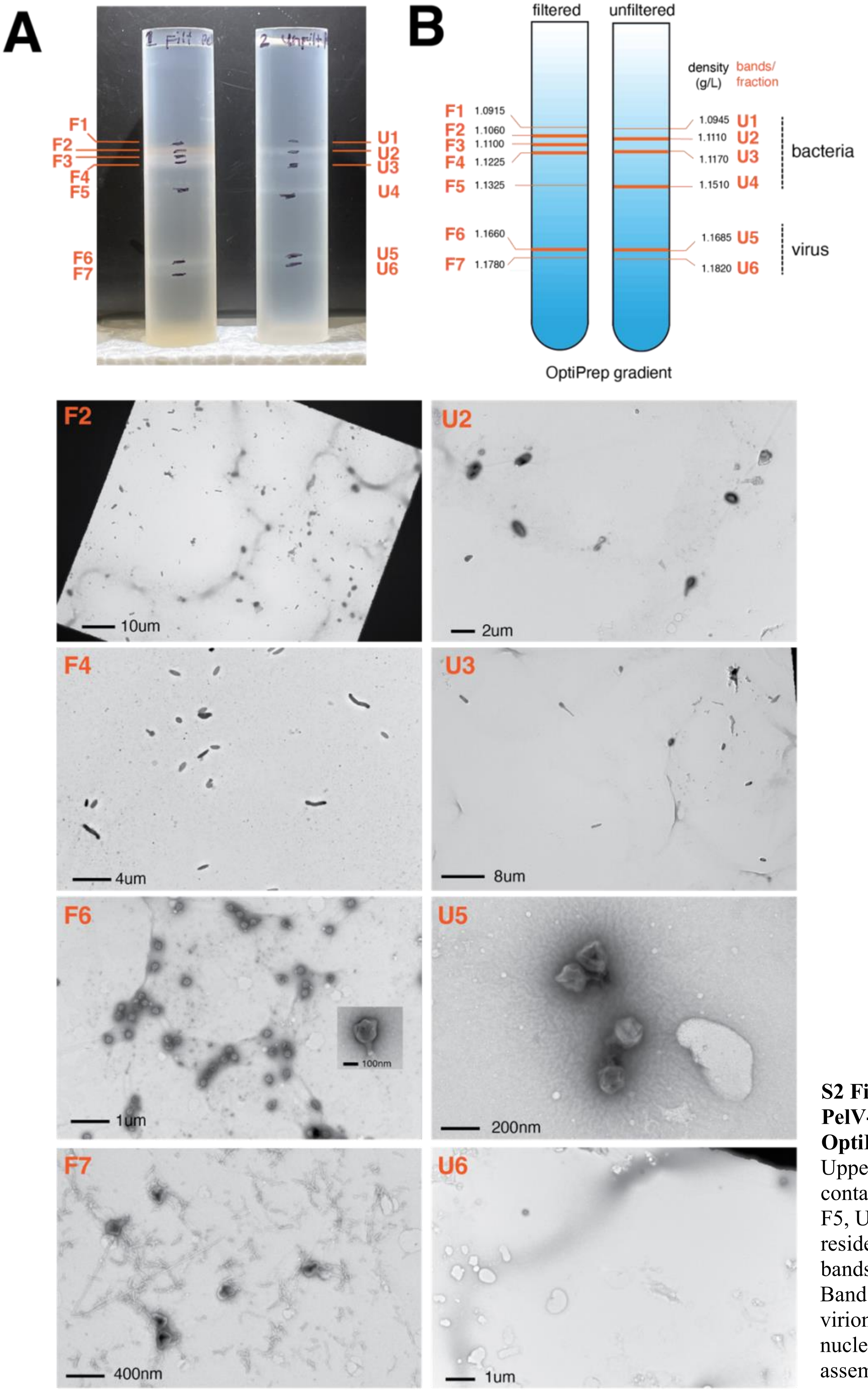


**S2 Figure. Purifying PelV-1 particles using an OptiPrep gradient.** Upper, less dense fractions contain bacteria (bands F1-F5, U1-U4), while viruses resided in more dense bands (F6-F7, U5-U6). Band F7, aside from virions, contains the nucleoprotein-like assemblies.

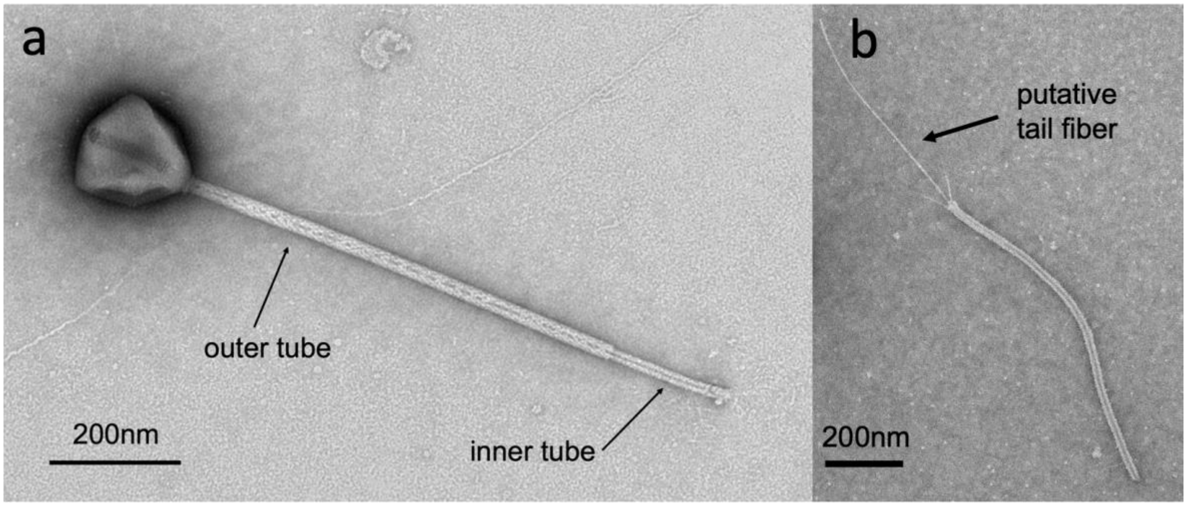


**S3 Figure. PelV-1 tail features.** (a) A typical tailed virion with outer and inner tubes. (b) In some cases, we observed detached inner tubes with putative tail fibers coming out of its end.

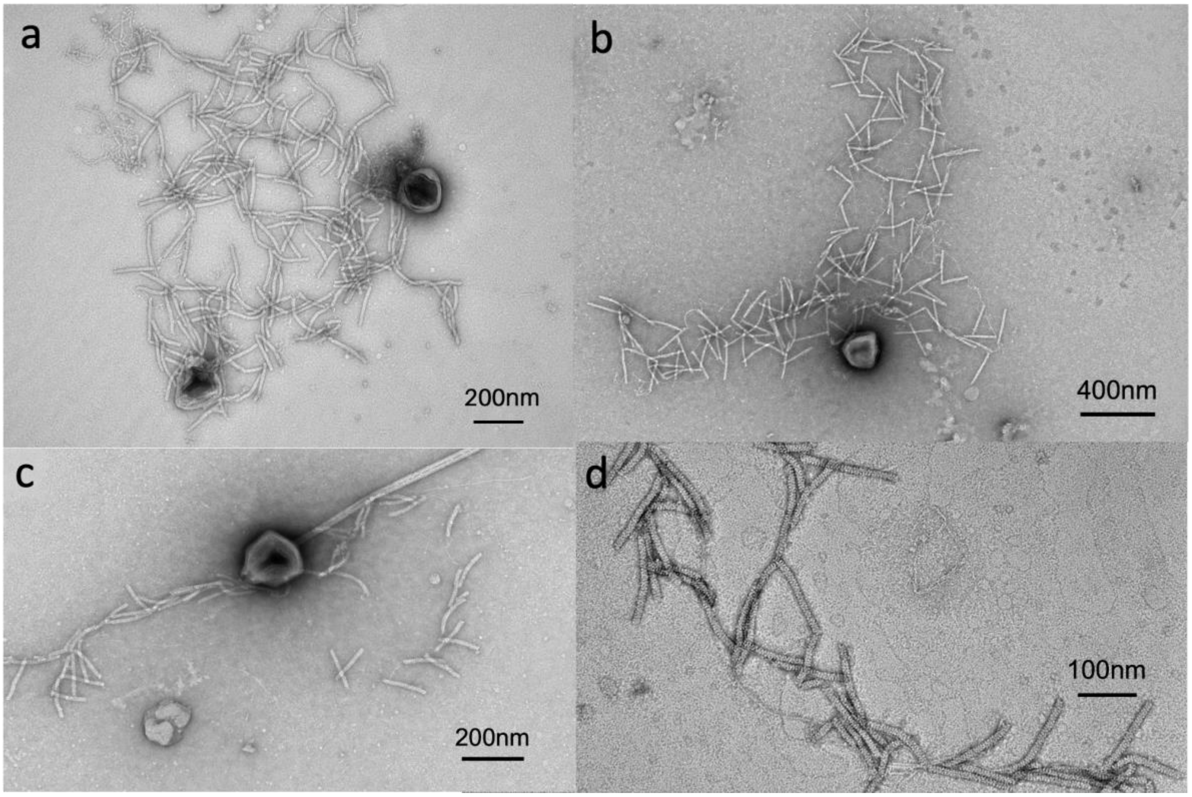


**S4 Figure. Nucleoprotein-like assembly (NLA)** (a-c). Fibrous structures are often found in the vicinity of the capsid which we call nucleoprotein-like assembly due to its morphological similarity with APMV nucleoprotein assembly/genomic fiber (Villalta et al., 2022) (d) A close-up of NLA shows a distinct appearance than tail structures.

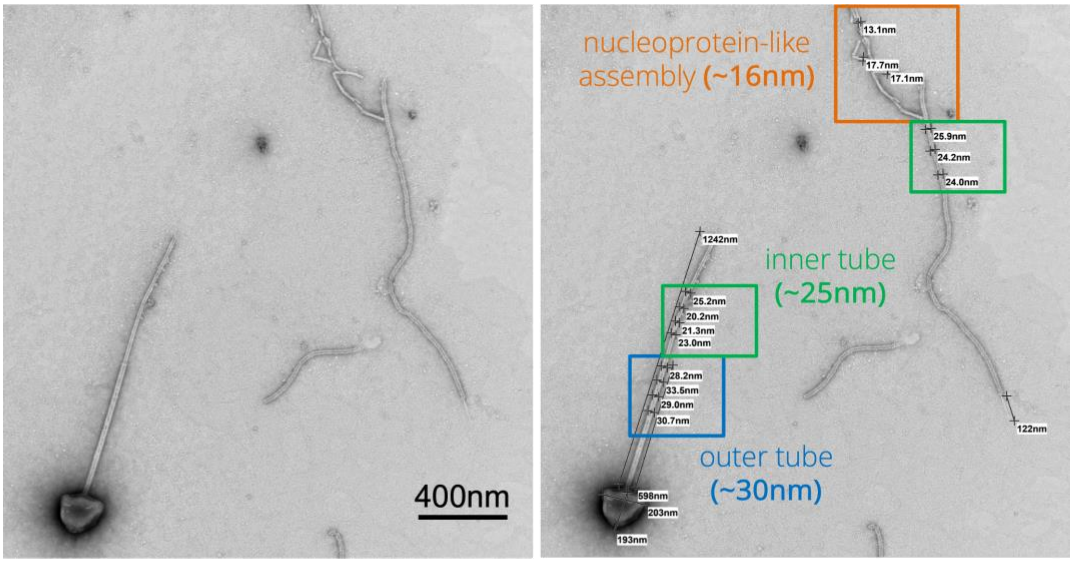


**S5 Figure. Comparison of tail and filamentous nucleoprotein-like assembly.** Thickness and surface appearance of virus tail (25-30 nm) and NLA (~16 nm) shows distinct structures.

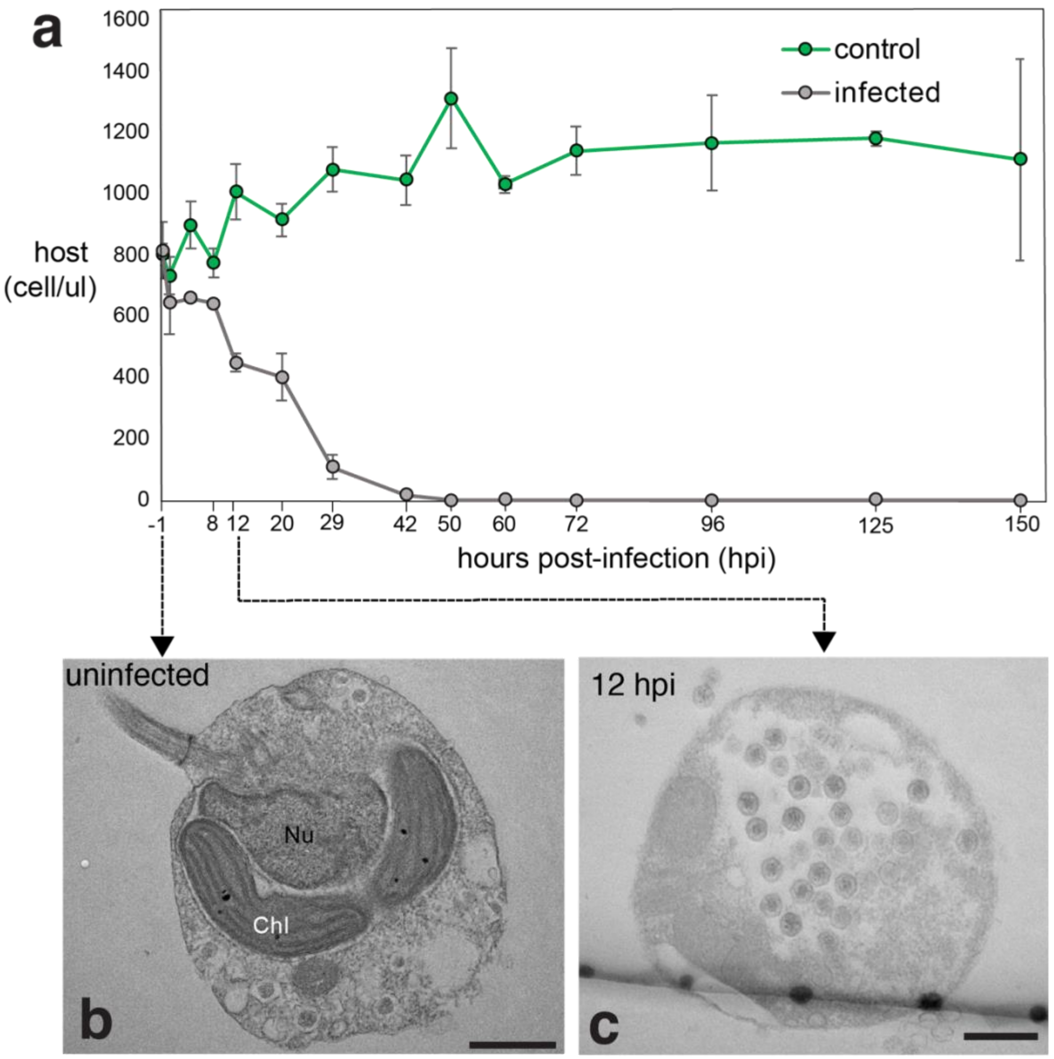


**S6 Figure. The PelV-1 infection time series (March 2022)** (a) Flow cytometry measurements of *Pelagodinum* (host) concentration across the infection time course. Concentration decreased at 8 hpi, eventually clearing at 42 hpi (b-c). Ultrathin sections showed a non-tailed PelV-1 variant at 12 hpi. (Nu-nucleus; Chl-chloroplast). Scale bar = 800nm

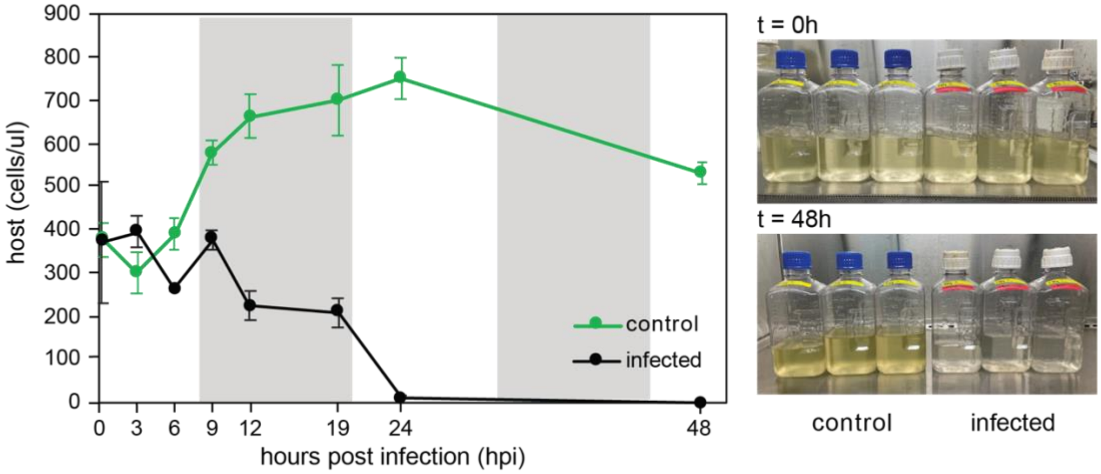


**S7 Figure. Infection time series (November 2022).** Healthy and infected cultures were monitored until 48 hpi. Samples were obtained for virus attachment and egress observations under TEM and SEM. Flow cytometry counts show a decrease in host concentration, which eventually cleared at 24 hpi.

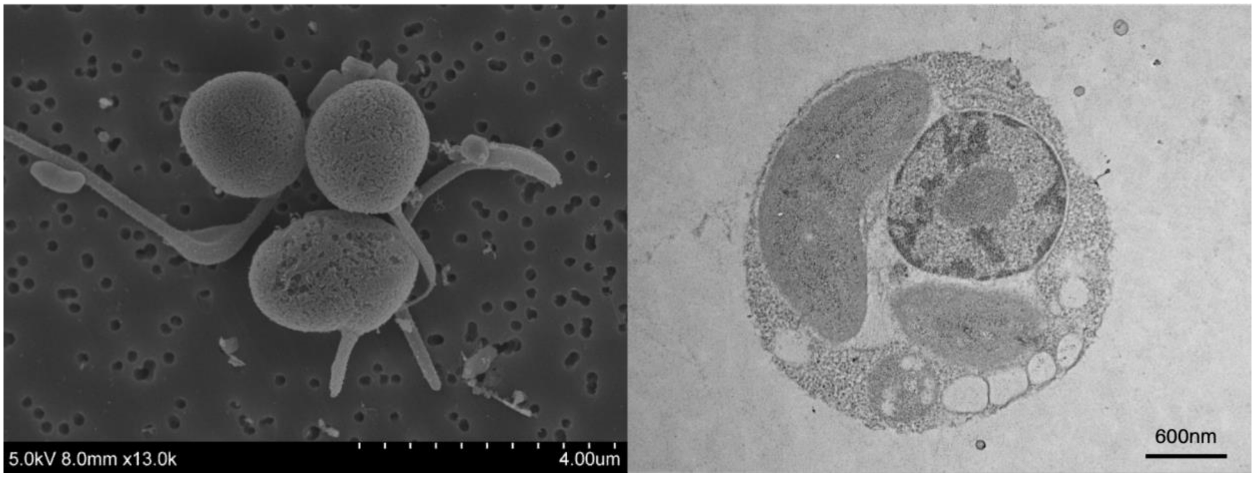


**S8 Figure. *Pelagodinium* sp. morphology and ultrastructure.** (A) Scanning electron micrographs showing the cell morphology and flagella. Typically, *Pelagodinium* is armored, but here, thecal plates are undiscernible, most likely because of the harsh post-fixation method. (B) Typical eukaryotic organelles, including the seemingly condensed chromosome typical of dinoflagellates, are shown.

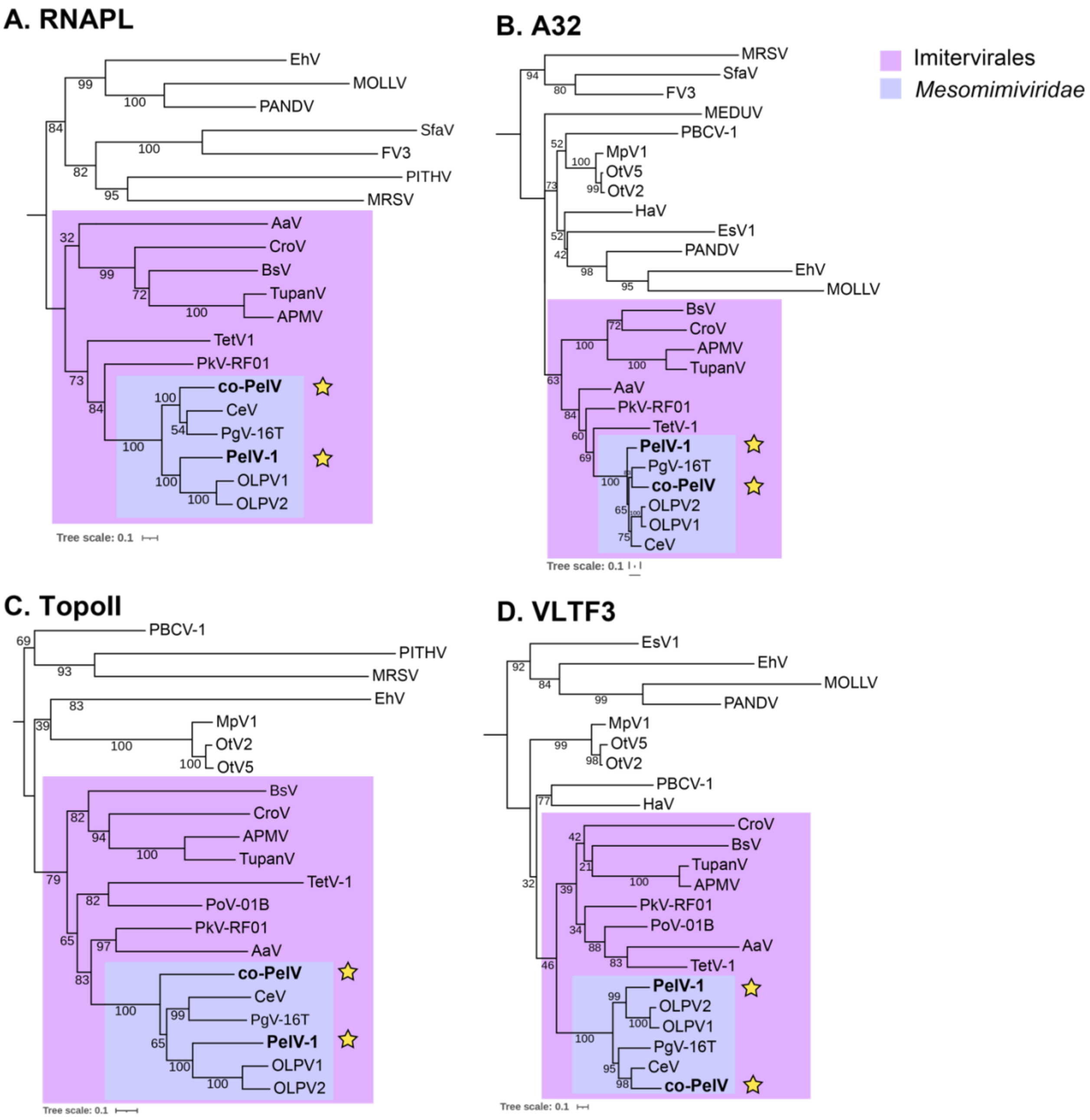


**S9 Figure. PelV-1 and co-PelV phylogenies based on multiple NCLDV markers** (RNAPL, A32, TopoII, and VLTF3). PelV-1 and co-PelV are placed among the Order Imitervirales, Family IM-01, Mesomimiviridae.

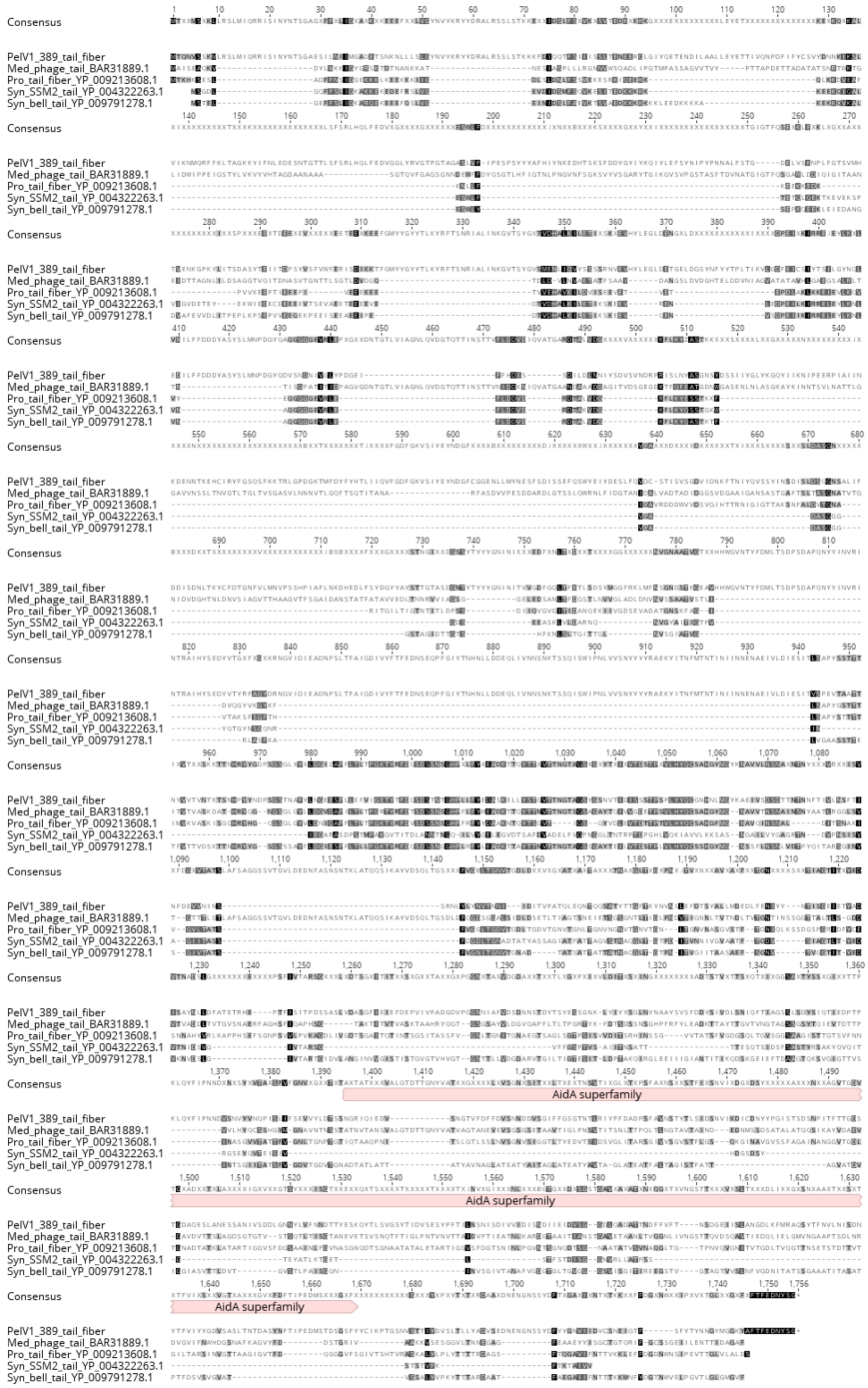


**S10 Figure. Protein alignment and location of AidA conserved domain across viral tail fiber protein.** Name and protein accession numbers are indicated. Alignment (using MUSCLE) and figure were made using Geneious 11.1. The conserved domain was identified using NCBI conserved domain search.

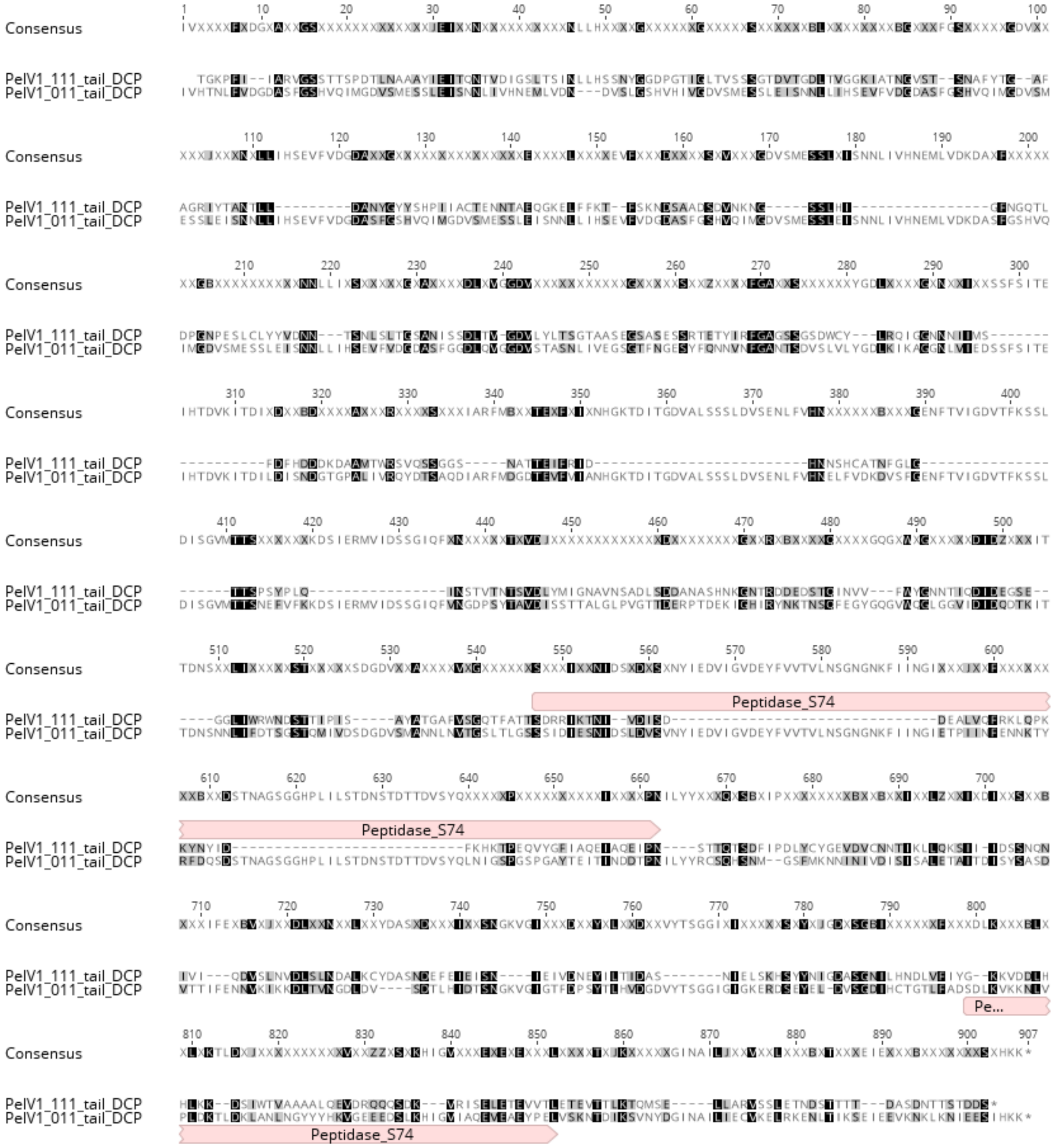


**S11 Figure. Protein alignment and location of Peptidase_S74 conserved domain (of bacteriophage tail protein) on the two other PelV-1 tail fiber protein homologs.** Alignment (using MUSCLE) and figure were made using Geneious 11.1. Conserved domain location was identified using NCBI conserved domain search.

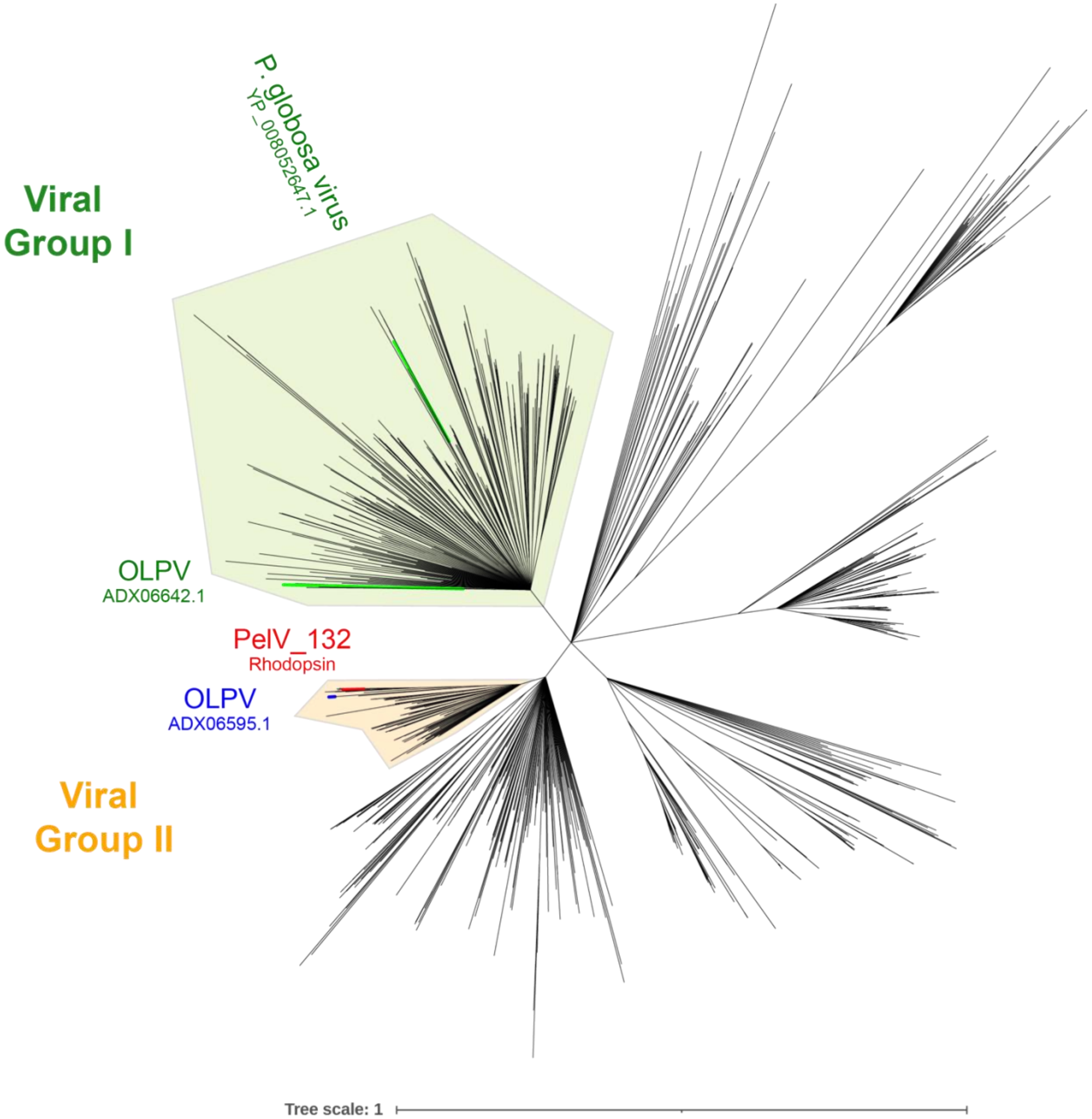


**S12 Figure. Phylogeny of PelV-1 rhodopsin.** Protein sequences from Schulz et al. 2020, including those derived from MAGs. Tree was made through MAFFT alignment, Trimal trimming and Geneious NJ tree builder. Visualized in iToL. Viral Groups 1 and 2 (VG1 and VG2) are noted based on Schulz et al 2020, highlighting OLPV and P. globosa virus as representative sequences. Other branches, not part of VG1 or VG2 represents mostly bacteriorhodopsin based on Schulz et al 2020.

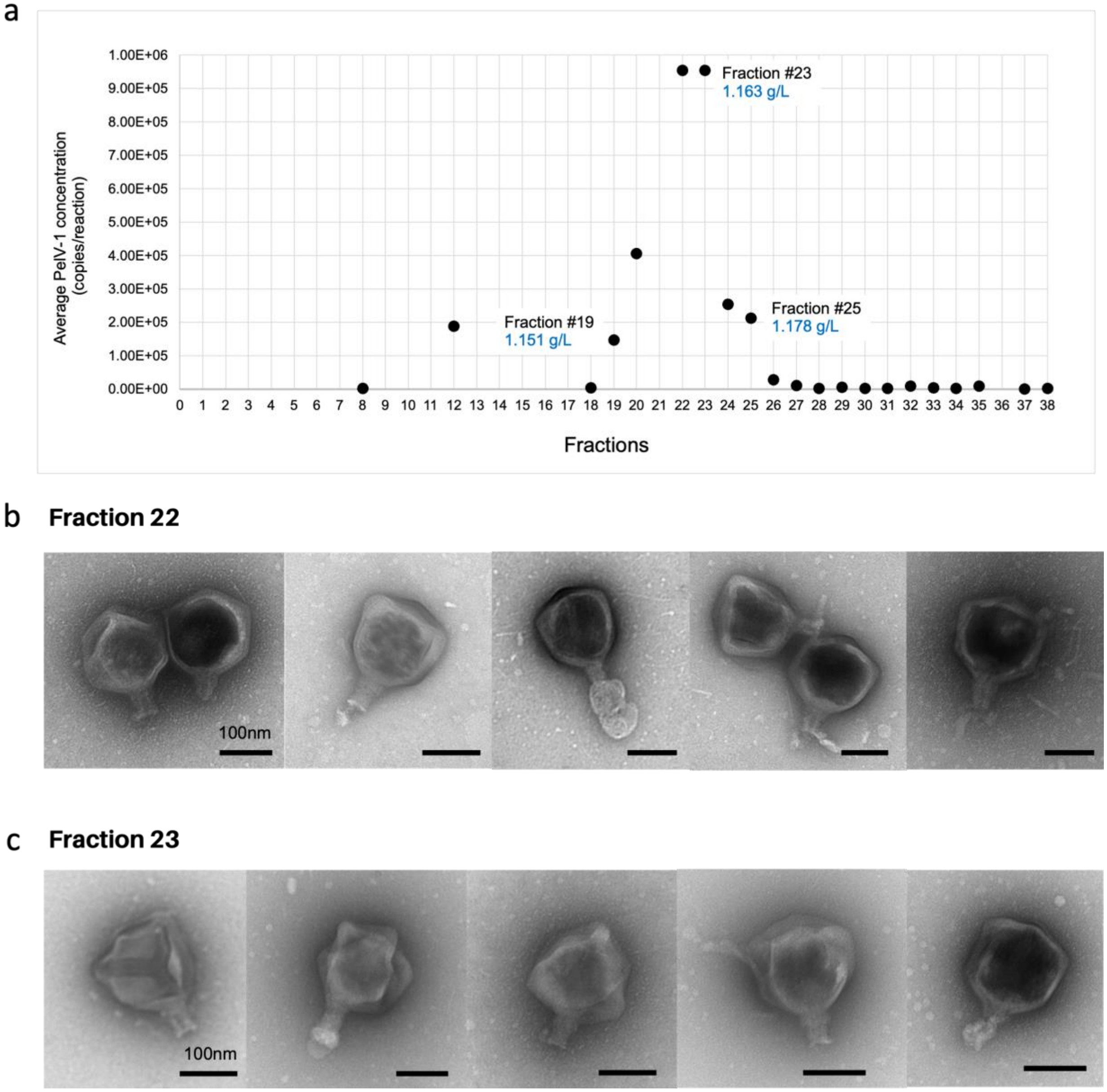


**S13 Figure. Confirming PelV-1 morphology** (a) An OptiPrep gradient consisting of 38 fractions were screened for PelV-1 abundance via QPCR. Fractions #22 and #23 were confirmed as the peak of PelV-1. (b-c) Representative virion morphology in fractions #22 and #23 confirmed the presence of either tail-like appendage or stubby protrusion. Appendage thickness ranges from 30 – 50 nm. Scale bar at 100nm.

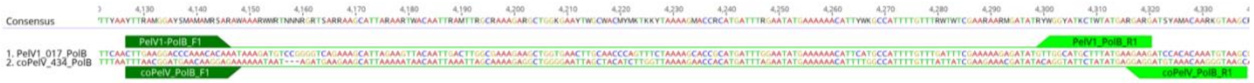


**S14 Figure: Alignment of primers to PolB.** PolB nucleotide sequences from both PelV-1 and co-PelV were aligned. Forward (in dark green) and reverse primers (in light green) were designed covering ~200bp region.

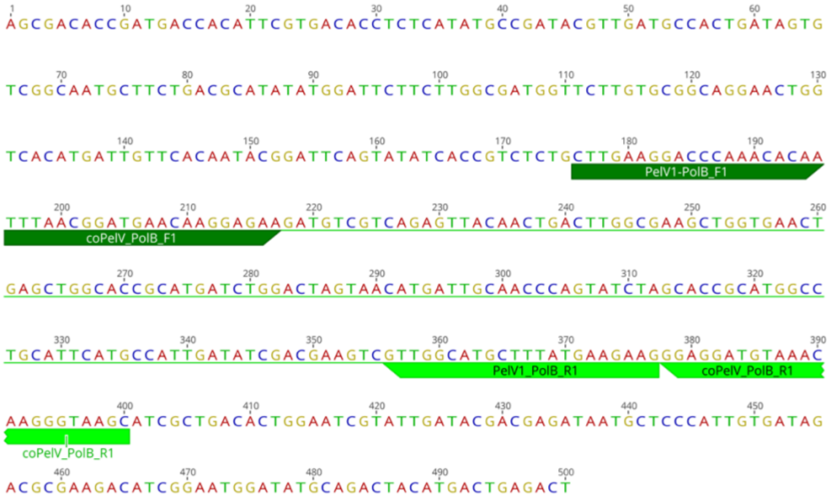


**S15 Figure. Alignment of primers to gBlock™**. A 500 nt gBlock was designed as reference for absolute quantification of PolB genes for both viruses.

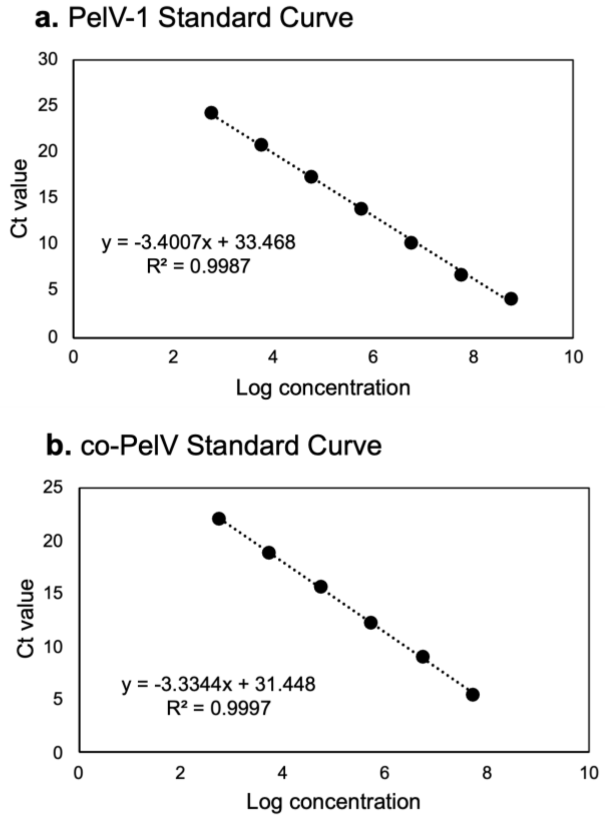


**S16 Figure. PelV-1 and co-PelV primer standard curve.** A dilution series was prepared using the gBlock™. Data is reported as average $C_t$ value of duplicate reactions.

**S1 Table.** Mean coverage of each bin

| bins | taxonomy | CsCl | | OptiPrep | | | |
|---|---|---|---|---|---|---|---|
| | | F11 | F17 | F6 | F7 | U5 | U6 |
| bin_1 | Unknown | 11.63 | 6.16 | 14.32 | 10.69 | 1.61 | 3.86 |
| bin_2 | Unknown | 0.18 | 0.03 | 4.90 | 5.96 | 0.10 | 0.51 |
| bin_3 | Bact_Lentilitoribacter | 0.54 | 0.26 | 11.68 | 3.97 | 2.52 | 4.44 |
| bin_4 | Bact_Shimia | 5.64 | 2.74 | 0.33 | 0.29 | 0.22 | 0.63 |
| bin_5 | Bact_H_grinnelliae | 2.54 | 0.59 | 8.45 | 10.62 | 6.54 | 13.89 |
| bin_6 | Unknown | 1.40 | 0.21 | 2.55 | 1.06 | 0.35 | 0.61 |
| bin_7 | Bact_M_mari | 0.94 | 5.77 | 0.07 | 0.06 | 0.25 | 0.77 |
| bin_8 | Bact_Alteromonas | 8.98 | 4.66 | 11.23 | 8.81 | 1.36 | 3.40 |
| bin_9 | Bact_Shimia | 22.37 | 1.13 | 2.51 | 1.55 | 2.56 | 4.35 |
| **bin_10** | **PelV-1** | 874.90 | 1892.64 | 3823.53 | 3681.30 | 3756.19 | 4218.39 |
| bin_11 | Bact_Balneola | 31.39 | 12.08 | 4.17 | 2.04 | 0.39 | 0.53 |
| **bin_12** | **co-PelV** | 0.42 | 2.71 | 0.25 | 24.12 | 0.15 | 7.85 |
| bin_13 | Bact_Muricauda | 20.11 | 1.71 | 4.22 | 2.21 | 0.11 | 0.23 |
| bin_14 | Bact_Hyphomonas | 22.61 | 2.35 | 15.32 | 6.80 | 7.45 | 7.27 |
| bin_15 | Bact_Pseudohongiellaceae | 1.86 | 2.79 | 0.44 | 0.15 | 0.59 | 1.79 |
| bin_16 | Bact_Marinobacter | 1.23 | 0.08 | 3.10 | 3.77 | 0.04 | 0.19 |

**S2 Table.** Percent (%) mapped reads to each bin.

| bins | taxonomy | CsCl fractions | | Optiprep fractions | | | |
|---|---|---|---|---|---|---|---|
| | | F11 | F17 | F6 | F7 | U5 | U6 |
| bin_1 | Unknown | 0.38 | 0.17 | 0.23 | 0.18 | 0.03 | 0.06 |
| bin_2 | Unknown | 0.00 | 0.00 | 0.03 | 0.03 | 0.00 | 0.00 |
| bin_3 | Bact_Lentilitoribacter | 0.19 | 0.08 | 2.00 | 0.73 | 0.48 | 0.73 |
| bin_4 | Bact_Shimia | 1.74 | 0.72 | 0.05 | 0.05 | 0.04 | 0.09 |
| bin_5 | Bact_H_grinnelliae | 1.25 | 0.25 | 2.04 | 2.75 | 1.77 | 3.22 |
| bin_6 | Unknown | 0.06 | 0.01 | 0.06 | 0.03 | 0.01 | 0.01 |
| bin_7 | Bact_M_mari | 0.15 | 0.80 | 0.01 | 0.00 | 0.02 | 0.06 |
| bin_8 | Bact_Alteromonas | 4.01 | 1.77 | 2.47 | 2.08 | 0.34 | 0.72 |
| bin_9 | Bact_Shimia | 8.68 | 0.37 | 0.48 | 0.32 | 0.55 | 0.80 |
| bin_10 | **PelV-1** | 37.52 | 69.06 | 80.73 | 83.32 | 89.01 | 85.35 |
| bin_11 | Bact_Balneola | 10.21 | 3.34 | 0.67 | 0.35 | 0.07 | 0.08 |
| bin_12 | **co-PelV** | 0.02 | 0.11 | 0.01 | 0.60 | 0.00 | 0.17 |
| bin_13 | Bact_Muricauda | 7.49 | 0.54 | 0.77 | 0.44 | 0.02 | 0.04 |
| bin_14 | Bact_Hyphomonas | 7.72 | 0.68 | 2.58 | 1.23 | 1.41 | 1.17 |
| bin_15 | Bact_Pseudohongiellaceae | 0.32 | 0.40 | 0.04 | 0.01 | 0.06 | 0.14 |
| bin_16 | Bact_Marinobacter | 0.41 | 0.02 | 0.51 | 0.67 | 0.01 | 0.03 |
| | **total** | 80.16 | 78.33 | 92.66 | 92.79 | 93.81 | 92.68 |

**S3 Table.** Number of reads belonging to each viral bin/genome

| | CsCl fractions | | OptiPrep fractions | | | | Raw lysate |
|---|---|---|---|---|---|---|---|
| | **F11** | **F17** | **F6** | **F7** | **U5** | **U6** | |
| **PelV-1 reads** | 2,675,713 | 5,788,281 | 11,693,540 | 11,258,558 | 11,487,593 | 12,901,169 | 5,124,909 (37.1%) |
| **co-PelV reads** | 1,401 | 9,122 | 853 | 81,089 | 515 | 26,377 | 70 (0.0005%) |
| **total reads** | 7,130,582 | 8,381,332 | 14,484,926 | 13,512,966 | 12,906,610 | 15,115,512 | 13,802,846 |

**S6 Table.** PelV-1 and co-PelV primers and gBlock™

| Target Gene | Primer | Product Length | Sequence (5‘ to 3‘) |
|---|---|---|---|
| PelV-1 polB | PelV1_PolB_F1 | 194 bp | CTTGAAGGACCCAAACACAA |
| | PelV1_PolB_R1 | | CTTCTTCATAAAGCATGCCAAC |
| co-PelV PolB | coPelV_PolB_F1 | 208 bp | TTTAACGGATGAACAAGGAGAA |
| | coPelV_PolB_R1 | | GCTTACCCTTGTTTACATCCTCC |
| gBlock™ (as reference) | | | AGCGACACCGATGACCACATTCGTGACACCTCTCATATGCCGATACGTTGATGCCACTGATAGTGTCGGCAATGCTTCTGACGCATATATGGATTCTTCTTGGCGATGGTTCTTGTGCGGCAGGAACTGGTCACATGATTGTTCACAATACGGATTCAGTATATCACCGTCTCTGCTTGAAGGACCCAAACACAATTTAACGGATGAACAAGGAGAAGATGTCGTCAGAGTTACAACTGACTTGGCGAAGCTGGTGAACTGAGCTGGCACCGCATGATCTGGACTAGTAACATGATTGCAACCCAGTATCTAGCACCGCATGGCCTGCATTCATGCCATTGATATCGACGAAGTCGTTGGCATGCTTTATGAAGAAGGGAGGATGTAAACAAGGGTAAGCATCGCTGACACTGGAATCGTATTGATACGACGAGATAATGCTCCCATTGTGATAGACGCGAAGACATCGGAATGGATATGCAGACTACATGACTGAGACT |
